# Integrated proteomic and phosphoproteomic profiling reveals mechanisms of Bisphenol-A induced placental toxicity

**DOI:** 10.64898/2026.03.04.709712

**Authors:** Ankit Biswas, Sandhini Saha, Naman Kharbanda, Tushar Kanti Maiti

## Abstract

The global industrialization and rapid urbanization elevated the risk of toxic pollutant exposure, which affects human health specially during pregnancy. Pregnant mothers are daily exposed to bisphenol-A (BPA), which is a common plastic leachate and a prominent toxic pollutant present in our environment. BPA act as an endocrine disrupting chemical (EDCs) by altering feto-placental homeostasis. This persistent and potent exposure of BPA during gestation can trigger placental damage affecting trophoblast cell function and survival. BPA even disrupts specific signalling cascades by altering post translational protein phosphorylation. However, this BPA mediated dysregulation of signalling nodes in early trimester placenta is still unexplored. Therefore, this study investigates the global proteome changes in post-BPA exposed extravillous trophoblast (EVTs) cells, which revealed a BPA mediated dynamic regulation of phosphoproteome-signatures and their associated kinases. Further inspection showed that the altered phosphorylation of c-JUN (S63) and GSK3α (Y279) is associated with BPA toxicity in EVTs and placenta. This altered phosphorylation affects the cellular signalling downstream, imparting damage upon the growing feto-placental unit. This highlights an altered phosphorylation mediated mechanism of BPA toxicity in placenta which can cause an onset of adverse pregnancy outcome. Data are available via ProteomeXchange with the identifiers PXD074780.

## 1. INTRODUCTION

A healthy pregnancy often requires a proper tuning of physiological homeostasis between the growing fetus and the mother. This physiological adaptation during pregnancy is supported by the placenta, facilitating the nutrient uptake, gas exchange and, waste excretion for the fetus^1^. However, these essential placental functions can be affected by various factors like maternal obesity^2^, nutritional stress^3^, immune dysregulation^4^ or environmental exposures to chemicals^5^. Due to rapid industrial growth, global urbanization and booming population, this toxic exposure increased many fold raising human health concerns, especially during pregnancy. Mothers during conception are usually exposed to such toxic chemicals like pesticides, heavy metals, air particulate matters or endocrine disrupting chemicals (EDCs) from plastics goods and everyday personal care products^6^.

Endocrine disrupting chemicals are natural or xeno-biotic compounds which can interfere with systemic hormonal homeostasis affecting development, reproduction and physiology. This EDCs can damage the developing feto-maternal axis due to their potent nature and persistent exposure during gestation^7^. Different classes of endocrine disrupting chemicals like phthalates, polychlorinated biphenyls, parabens and bisphenols are associated with various reproductive disorders^8^. However, bisphenols emerged as a model endocrine disruptor to study reproductive toxicity due to its widespread environmental exposure showing lower dose impact against broader range of cellular receptors^9^. Urine sampling of pregnant mothers across all trimesters detected low concentration of bisphenols, among which bisphenol-A (BPA) is present predominantly^10^. Previous studies highlighted the association of prenatal BPA exposure with preterm birth^11,12^ and fetal growth restrictions^13^. Whereas higher BPA concentration in maternal serum and urine from mid-gestation is associated with the onset of pre-eclampsia^14–16^. Similarly, experimentally impregnated mice models administered with different concentration of BPA can develop either fetal growth restriction^17,18^ or pre-eclampsia-like phenotype^19,20^. Thus, maternal exposure of bisphenol-A can elicits detrimental damage to feto-maternal axis ultimately contributing towards adverse pregnancy outcome. But the holistic mechanism of BPA mediated placental toxicity is still under-studied.

Bisphenol-A, absorbed from environmental sources in maternal circulation, can easily cross the placental barrier to get accumulated in fetal and placental tissues^21,22^ and then imparts adverse effect on the fetus altering organ development, fetal metabolism and growth^23^. This BPA-mediated adverse effect resulted from trophoblast cell injury, which disrupted placental homeostasis by inducing oxidative stress^24^, deregulating trophoblast gene expression^25^, and inhibiting trophoblast cell migration and invasion^26,27^. Similarly, BPA exposed murine placentas shows metabolic alteration^28^, impaired spiral artery remodeling^17^, epigenetic reprogramming^19^ and disorganization of placental layer morphology^29^. BPA also damages placenta-brain^30^ and gut-placental^31^ axis by affecting systemic regulation.

Protein tyrosine phosphorylation is elevated by BPA mediated activation of protein kinase A in rat sperm^32^ while BPA exposed human sperm shows reduced protein phosphorylation in tyrosine residue^33^. Similarly, BPA treatment on one hand, can impair neuronal insulin signalling by decreasing phosphorylation of IR, IRS1, GSK3B, AKT and ERK1/2^34^, while on other it can target AMPKα phosphorylation inducing neuronal apoptosis and degeneration^35^. Neuronal differentiation can also be suppressed by Bisphenol-A through reduction of ERK1/2 and MAPK phosphorylation^36^. Phosphorylation act as molecular switch, essential for signal transduction, metabolic regulation and energy production. This phosphorylation switch is often balanced delicately by regulation of kinase and phosphatase activity. However, there is a large knowledge gap surrounding the fate of phosphorylated proteins in placenta and trophoblast post-BPA exposure. Thus, we hypothesized that fatal exposure of bisphenol-A can alters the global phosphorylated protein pool in both trophoblast cells and placenta. Although, several studies revealed BPA mediated alteration of phosphorylation signatures in sperm and neurons can cause dynamic regulation of downstream signalling, but a placenta-centric comprehensive screening of BPA regulated phospho-signatures is essential for deciphering the mechanism of BPA mediated placental toxicity.

Therefore, this current study explores the BPA altered phosphoprotein dynamics in extravillous trophoblast cells which is further validated in placental tissues post BPA exposure. We integrated a differential proteome with a dynamic phospho-proteome of early trimester trophoblast cells after BPA treatment. Our study revealed a bisphenol-A mediated kinase regulation, which altered the trophoblast phosphoprotein signatures and triggers dysfunction of cellular signalling essential for healthy functioning of placenta. Furthermore, the present study also showed BPA-mediated differential phosphorylation of GSK3A and c-JUN in trophoblast cells, which was similarly observed in placentas from a BPA-treated murine model. This highlights BPA induced alteration of phosphoprotein-dynamics, as a molecular basis for bisphenol-A mediated placental toxicity during pregnancy.

## 2. EXPERIMENTAL PROCEDURE

### 2.1. Cell culture and treatment

First-trimester human placenta derived immortalized extravillous trophoblast cells (HTR8/SVneo) were maintained in Roswell Park Memorial Institute (RPMI) 1640 medium (R6504, Sigma) containing 2g/L sodium bicarbonate (40151UR-K05, SDFCL), 100 units/mL penicillin, 100 µg/mL streptomycin (A001A, Himedia) and supplemented with 10% heat-inactivated fetal bovine serum (FBS) (A5256501, Gibco). The cells were kept under humidified conditions at 37 °C in 5% CO2. 3X10^5^ cells per well were seeded in a 6 well plate for experiments after passaging for at least 3 times from revival. The cells were grown overnight until approximately 80% confluent and then serum-starved with 1% FBS supplemented incomplete RPMI-1640 media for 4 hours. Then the cells were subsequently treated with final concentration of 10 µM bisphenol-A (239658, Sigma) dissolved in DMSO (472301, Sigma) for 24 hours. The bisphenol-A concentration was pre-determined using assays in earlier study. Cells treated with 0.1% DMSO for 24 hours were considered as mock control.

### 2.2. Global differential proteomics

#### 2.2.1. Sample preparation and Tryptic Digestion

The differential proteomics analysis was performed after cells were treated with 10 µM BPA or mock control as described earlier. Briefly, 3X10^5^ cells/well were seeded in a six well plate and then serum-starved for 4 hours once it reaches 80% confluency. Then, the respective wells were exposed to BPA or kept as mock control for 24 hours and then harvested in RIPA (R0278, Sigma) lysis buffer containing 1X halt-protease and phosphatase inhibitor cocktail (78441, Thermo Scientific). The lysates were homogenized and probe sonicated to ensure optimum extraction of total protein. The extracted proteins were then precipitated using chilled acetone (014-08681, Wako). The precipitates were centrifuged, air-dried and redissolved in 8M Urea (219-00175, Wako). The proteins dissolved in 8M urea were diluted till 2M urea concentration using 100 mM ammonium bicarbonate (ABC) (A6141, Sigma) buffer and protein concentration was quantified using BCA protein assay kit (23225, Pierce, Thermo Scientific).

Equal amount of protein (50 µg) was taken from each sample and performed reduction using 10 mM DL-Dithiothreitol (DTT) (D5545, Sigma) at 56 °C for 45 mins followed by alkylation with 20 mM Iodoacetamide (IAA) (I1149, Sigma) for 1 hour at room temperature. Next, the reduced and alkylated samples were incubated using 1:20 (enzyme: protein) ratio of MS grade trypsin (90057, Pierce, Thermo Scientific) for overnight at 37 °C. After, overnight incubation the digestion was quenched by drying the samples using vacuum-centrifuge. The samples were then reconstituted and passed through Oasis HLB C18 cartridges (WAT094225, Waters) using an Extraction Manifold (WAT200607, Waters, USA) for desalting. The desalted tryptic peptides were again vacuum dried and stored until ready for LC-MS/MS analysis.

#### 2.2.2. Data acquisition and protein identification

The dried and desalted tryptic peptides were reconstituted in solvent A (2% [vol/vol] acetonitrile, 0.1% [vol/vol] formic acid in water) for LC-MS/MS analysis. The data were acquired in Zeno-SWATH mode using a ZenoTOF 7600 MS (Sciex) linked with a waters microscale LC system (ACQUITY UPLC M-Class System). 500 ng tryptic digest from each sample were loaded onto Luna 5 µm C18 Micro trap column (20 mm X 0.3 mm, 100 Å, Phenomenex) and then resolved using a nanoEase M/Z HSS T3 analytical column (150 mm X 300 µm, 1.8 µm, 100 Å, Waters) with a flow rate of 5 µL/min. The peptides were separated utilizing a linear gradient of solvent B (100% (v/v) acetonitrile with 0.1% (v/v) formic acid) for 22 minutes, with final run duration of 32 minutes. 65 variable sized overlapping windows were used to scan MS2 mass ranged from 350-950 Da utilizing 20 msec accumulation window and 1.74 sec per cycle period for a total 1103 cycles during a single run.

The SWATH acquired .wiff files were utilized to build de-novo spectral library against the UniProtKB human protein database with isoforms (42,502 entries, December 2024) in Spectronaut pulsar 19 (Biognosys) employing default parameters. The SWATH-MS data were then searched against the generated spectral library with standard search settings: for RT prediction dynamic iRT was utilized, interference correction was enabled at MS2 level, and data normalization was done by global TIC (total intensity count). A strict 1% FDR was used for both precursor and peptide filtering ensuring maximum possible true detection for proteins. Finally, trypsin is selected for specific enzyme allowing maximum two missed cleavages. Carbamidomethyl on cysteine (+57.021464 Da) is used as fixed modification whereas, oxidation of methionine (+15.994915 Da), and N-terminal acetylation (+42.010565 Da) is considered as variable modifications.

The extracted peptide ion peak intensities for corresponding proteins were exported as PGquant values along with fold change and significance, which were ultimately analyzed in R software (R 4.3.1). The differential proteins were used for gene set enrichment analysis (GSEA) using ClusterProfiler (v4.12.6) against ReactomePA (1.48.0) database. The gene and pathway interaction map was plotted using cnetplot function in ClusterProfiler (v4.12.6) package.

### 2.3. Global phospho-proteome profiling

#### 2.3.1. Sample preparation and tryptic digestion

The phospho-proteome profiling was performed utilizing metal oxide based phospho-peptides enrichment in tandem with high resolution LC-MS/MS analysis. 3 X 10^6^ HTR8/SVneo cells were seeded in each 100 mm plates and incubated until the confluency reached 80%. The cells were exposed either with 10 µM BPA or kept as control for 24 hours and then collected in RIPA (R0278, Sigma) buffer containing 1X halt-protease and phosphatase inhibitor cocktail (78441, Thermo Scientific). Three 100 mm plates were pooled for each replicate to achieve sufficient protein amount for phospho-peptide enrichment. The harvested cells were homogenized, and probe sonicated to extract the total protein that was precipitated further with six volumes of chilled acetone. The acetone precipitated pellets were air-dried and redissolved in 8M Urea which were then diluted to 2M urea using 100 mM ABC buffer. Finally, the protein concentration was measured using BCA protein assay kit (23225, Pierce, Thermo Scientific) and 10 mg protein from each replicate were taken for next steps.

Sample reduction was performed by adding 10 mM DTT to the samples and incubating for 1 hour at 56 °C. The reduced proteins were then alkylated with 20 mM IAA for 1 hour at room temperature. Finally, MS grade trypsin (T1426, Sigma) was used in 1:10 (enzyme: substrate) ratio for overnight trypsin digestion of the samples at 37 °C. Following digestion, the reaction was terminated by acidification with 1% formic acid (FA) (065-04191, Wako). The peptide mixture was then centrifuged at 14,000g for 30 minutes to remove the undigested fractions. The supernatant was loaded onto reversed-phase C18 Sep-Pak cartridges (Cat no. WAT020515, Waters, USA) for peptide cleaning and desalting. The desalted peptides were then vacuum-dried and stored at -80 °C until enrichment.

#### 2.3.2. Phospho-peptide enrichment

The phospho-peptide enrichment was carried out using Titansphere titanium dioxide (TiO_2_) beads (5020-75010, GL Sciences) as mentioned in the earlier study^37^. Briefly, the TiO_2_ beads were equilibrated in 80% acetonitrile + 1% FA dissolved 2,5-dihydroxybenzoic acid (20 mg/ml) (149357, Sigma) for 30 minutes. Next, the vacuum-dried peptides were reconstituted in 10 ml of 80% acetonitrile + 1% FA followed by 30 minutes incubation with equilibrated TiO_2_ beads while rotating. Post incubation the samples were centrifuged at 4000X g for 5 minutes to remove the supernatant. The beads were thoroughly washed three times with sequentially higher concentrations of acetonitrile (30%, 50% and 80%) in 1% FA. Further, the phospho-peptides were sequentially eluted in 100 µl of 5% ammonium hydroxide (NH_4_OH) (40205, Sigma) and then in 100 µl of 10% NH_4_OH with 25% acetonitrile. The eluted phospho-peptides were vacuum-dried and desalted with C18 tips (Pierce, Thermo Fisher Scientific, USA) followed by reconstitution in solvent A (2% [vol/vol] acetonitrile, 0.1% [vol/vol] formic acid in water) for LC-MS/MS characterization.

#### 2.3.3. Mass-spectrometric analysis of enriched phospho-peptides

The LC-MS/MS analysis was performed utilizing ZenoTOF 7600 MS (Sciex) linked with a waters microscale LC system (ACQUITY UPLC M-Class System). Equal amount of tryptic peptides (1 µg) from each samples were injected in triplicates with a flow rate of 5 µL/min onto Luna 5 µm C18 Micro trap column (20 mm X 0.3 mm, 100 Å, Phenomenex) and then separated using a nanoEase M/Z HSS T3 analytical column (150 mm X 300 µm, 1.8 µm, 100 Å, Waters,) over a linear gradient of solvent B (100% (v/v) acetonitrile with 0.1% (v/v) formic acid) for 37 minutes, followed by washing and equilibration with solvent A for the remaining of 50 minutes. The MS data was acquired in IDA (information-dependent acquisition) mode utilizing OptiFlow Turbo V ion source controlled by SciexOS 3.1 with optimized settings for voltage, curtain gas, nebulizer gas, heater gas, and the source temperature. A single IDA cycle is consisted of survey (MS1) scan (m/z 300-1500 Da) of ions with charge state ranging from +2 to +5 for 100 msec, followed by a scan of top 45 MS/MS (200-1800 m/z) ions each for 20 msec, accumulating a total 1.24 sec for each cycle period. In the initial survey (MS1) scan a minimum tolerance of 30 ppm and, a dynamic exclusion window of 5 sec was applied for precursors with intensity above 100 cps. A dynamic collision energy with spread 5, and de-clustering potential of 80 were used for both MS1 and MS2 fragmentation.

#### 2.3.4. Phospho-peptides identification

The IDA acquired raw MS data were converted into mzML format utilizing MSconvertGUI (v3.0) from Protewizard^38^ with vendor peak picking. The converted mzML files were subsequently searched using MSFragger (v4.4.1)^39^ in Fragpipe (v23.0) against the UniProtKB human protein database with isoforms (42,502 entries, December 2024). Reversed protein sequences were incorporated in the protein database as decoys. The default LFQ-phospho workflow was followed for searching the mock and BPA group separately to identify unique protein signature in each condition. The tolerance for precursor and fragments (initial) was set to 20 ppm with mass calibration and parameter optimization applied. Isotope error was kept at 0/1/2/3, while enzyme specificity was selected as “strict trypsin” with maximum missed cleavage of 2. Peptides of 7 to 50 amino acids length and 500 to 5000 Da mass range were filtered for next step. Methionine oxidation (+15.9949, maximum 3 occurrences), protein N-terminal acetylation (+42.0106, maximum 1 occurrence) and, S/T/Y phosphorylation (+79.96633, maximum 3 occurrences) were used in variable modification. While, in fixed modification cysteine carbamidomethylation (+57.02146) was selected. Maximum 3 modifications per peptide was allowed. The MSFragger identified MS/MS spectral matches were re-scored using deep learning algorithm of MSbooster^40^ (v1.3.9) to generate peptide spectrum matches (PSMs). Percolator (v3.7.1) validates this PSMs for modification site identification by PTMprophet (v3.0.1). The results from Percolator and PTMprophet were processed by Philosopher^41^ (v5.1.1) for filtering of PSM, peptides, and proteins with 1% false discovery rate (FDR) cutoff. IonQuant^42^ (v1.11.9) performs MS1-based label free quantitation using this validated, high confidence peptide and protein abundance across the samples, with match between run (MBR) algorithm enabled.

After successful identification of phospho-sites and their associated proteins in BPA and mock condition separately, we overlapped them using Venny (v2.1) to segregate unique and common phospho-site and phospho-proteins between the two conditions. The pathway enrichment of unique and common proteins was performed using curated Reactome database in ShinyGO^43^ (v0.77) (https://bioinformatics.sdstate.edu/go77/) platform. The data were plotted using SRplot^44^ (https://www.bioinformatics.com.cn/en).

### 2.4. Animal and treatment

The animal experiments were approved by the Regional Centre for Biotechnology Institutional Animal Ethics Committee (IAEC Project No. RCB/IAEC/2021/092). Based on the approved animal use proposal eight female and four male C57BL6 mice (five to six weeks old) weighing 19-25 grams were issued from experimental animal facility (EAF), Regional Centre for Biotechnology. The mice were housed in polypropylene cages with twelve hours of light-dark cycles and ambient temperature under controlled environmental conditions. The mice were provided with standard chow diet and sterilized water in glass bottle ad libitum. A week of habituation period was given to the females before treatments were initiated. Eight female mice were randomly distributed in two groups. First group received oral gavage of bisphenol-A (200 µg/kg body weight per day dissolved in 100% ethanol and administered with corn oil for better adsorption). The mock control group were gavaged 100% ethanol in corn oil. The BPA dose was selected based on previous study^30^ where the concentration used was below the diet-administered maximum nontoxic dose for rodents (200 mg/kg body weight per day)^45,46^. Each female mice received the gavage daily for two weeks according to their respective group and subsequently paired with the potential breeder male in 2:1 ratio. Next morning the vaginal plugs were checked to determine successful mating. The presence of vaginal plug was inferred as e0.5 post-coitus. If the vaginal plug was absent in the morning, then the males were separated and repaired again that evening. The females were continued with their respective treatment till e12.5 when they were humanely euthanized to surgically remove the uterine horn containing the embryos. The embryos were carefully incised to harvest the placenta which were subsequently flash freezed in liquid N_2_ and stored at -80 °C.

### 2.5. Western blot analysis

The placental tissues and cellular samples from both treatment and control group were homogenized using radio immunoprecipitation assay (RIPA) lysis and extraction buffer (R0278, Sigma) containing 1X halt-protease phosphatase inhibitor (78441, Thermo Scientific). The lysate was probe sonicated to ensure maximum protein extraction followed by centrifugation at 4 °C. The supernatant containing total extracted proteins were quantified using BCA protein assay kit (23225, Pierce, Thermo Scientific). 40 µg protein from each sample were loaded in 10% sodium dodecyl sulphate-polyacrylamide gel (SDS-PAGE) gel for electrophoretic separation and then subsequently transferred to a 0.45 µ polyvinylidene fluoride (PVDF) membrane (IPVH00010, Millipore, Merk). After 1 hour blocking in 5% (W/V) skimmed milk (GRM1254, Himedia) the respective blots were incubated overnight at 4 °C with primary antibodies against GSK3α (1:2000, A19060, ABclonal), phospho-GSK3α-Y279 (1:2000, AP0261, ABclonal), c-JUN (1:1000, 9165, CST), phospho-c-JUN-S63 (1:1000, 2361, CST), PBK (1:1000, 4942, CST) and GAPDH (1:10000, A19056, ABclonal). Following primary antibody incubation, the blots were washed with Tris-buffered Saline-Tween (TBST) (0.1% [vol/vol]) and further incubated with anti-rabbit HRP-conjugated secondary antibody (1:10000) for 1 hour at room temperature. After secondary incubation the blots were washed with TBST and developed using Immobilon Forte Western HRP Substrate (WBLUF, Millipore, Merk). Digital images were captured using Image Quant LAS 4000 (GE Healthcare Bio-Sciences AB, Sweden) and subsequently quantified using Fiji (v2.17.0) software. The relative intensity of each protein was normalized against GAPDH. While the phospho-proteins were normalized using the total protein intensity of the same non-phosphorylated form.

### 2.6. Identification of kinase-substrate network

The list of known human kinases was downloaded from KinMap^47^ (http://www.kinhub.org/kinases.html) and overlapped with the differential proteome data to identify 75 kinases. After, q-value filtering, the kinases with their log2 fold change (FC) were mapped onto human kinome tree using CORAL^48^ online tool (http://phanstiel-lab.med.unc.edu/CORAL/). The identified kinases were enriched for pathways using KEGG database in ShinyGO (v0.77). Further, the list of phospho-sites targeted by the significantly regulated kinases were exported from PhosphoSitePlus^49^ along with their conserved motif diagrams (https://www.phosphosite.org/homeAction.action).

### 2.7. Transcription factor target prediction

The predicted list of c-JUN target proteins in both human and mice database were extracted from TFLink^50^ (https://tflink.net/) webtool. The list was filtered for targets identified only by small scale evidence to ensure confident prediction. Pathway enrichment of the filtered targets was done using curated Reactome database in ShinyGO (v0.77) based on their respective ontology.

### 2.8. Data visualization and statistical analysis

The data were plotted and visualized using Prism 8 software (GraphPad Software, USA) or ggplot (3.5.1) package in R(v4.3.1) if not either mentioned. The heatmaps and upset plots were generated by pheatmap (1.0.13) and UpsetR (1.4.0) package respectively. All the graphical illustrations were created using BioRender (https://www.biorender.com/).

Statistical analysis was performed using Prism 8 for all the experiments. The data-points were plotted at least for three biological replicates and expressed as mean ± Standard error of mean (SEM). The significance between the difference of control and BPA groups were determined using student’s t-test where statistical significance was considered for P values *<* 0.05 and are indicated by asterisks as follows: *P *<* 0.05, **P *<* 0.01, ***P *<* 0.001, ****P *<* 0.0001.

## 3. RESULTS

### 3.1. Bisphenol-A alters trophoblast proteome dynamics

Bisphenol-A (BPA) exposure to cells of placental lineages like HTR8/SVneo^27^, BeWo^24^ or JEG-3^51^ can cause detrimental effect on specific cellular nodes, as suggested by previous literature. However, the global effect of BPA exposure to this placental cell lines are poorly understood. Thus, we used HTR8/SVneo cells as a model for studying BPA toxicity in placenta. The 24-hour exposure of 10 μM BPA was selected based on both literature and previously performed assays. **Figure 1A** explains the general schema of the ZenoSWATH proteomics workflow followed in this study. The data distribution and quality were explored using pair plot (**Figure 1B**) where the lower off-diagonal panels display the pairwise scatter plots. The points representing protein intensities from paired samples, lies near to the red regression line suggestive of a good correlation between the samples. This is also evident from significant Pearson’s correlation coefficient presented on the upper off-diagonal panels for pairwise runs. The diagonal plots, showing protein intensity histogram for each replicate in both conditions, followed a Gaussian distribution demonstrating a good data quality sufficient for proceeding to next step.

**Figure 1:**
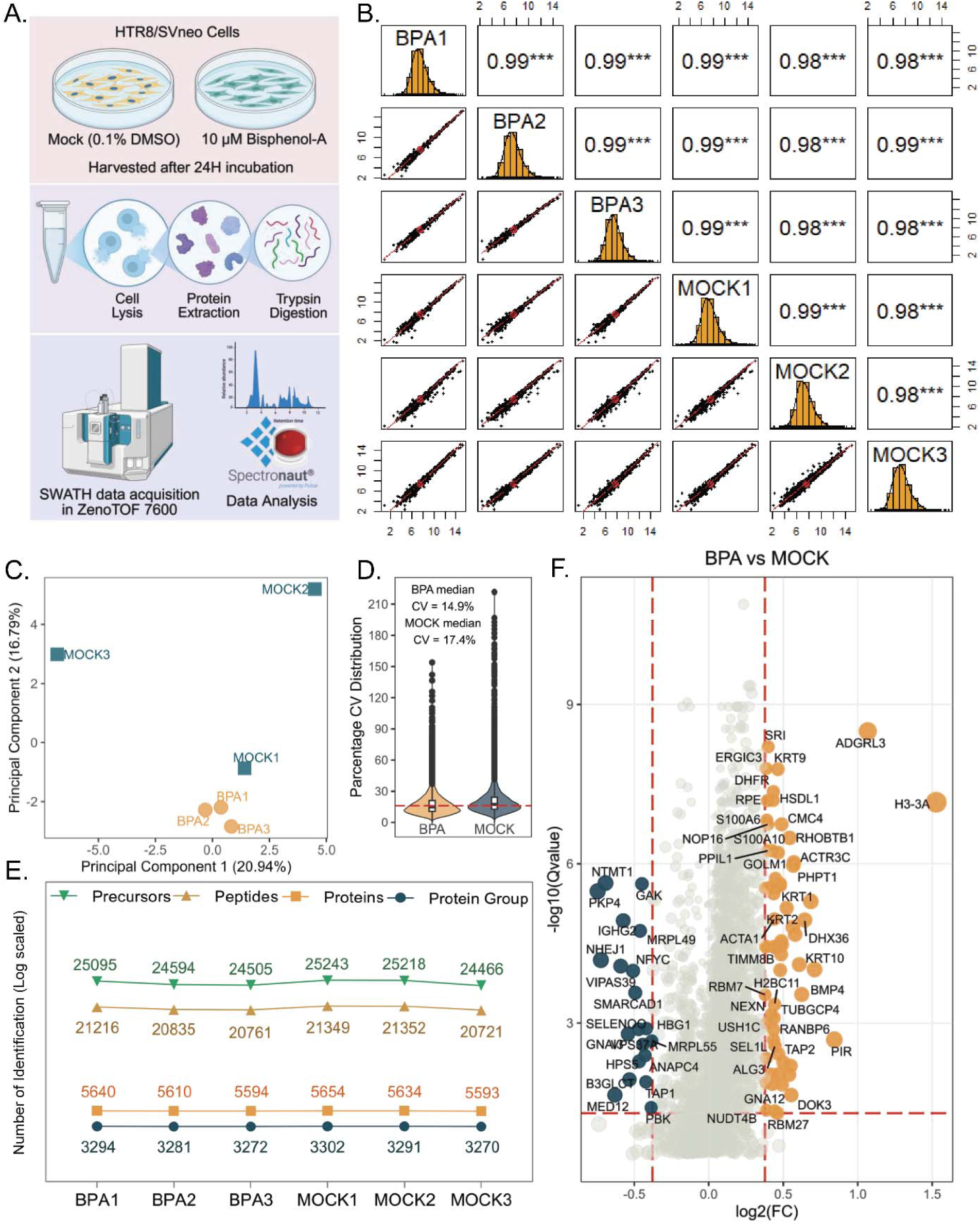
First trimester extravillous trophoblast cell proteome quality and differential pattern. **A)** The overview of Zeno-SWATH workflow followed for studying differential proteome. **B)** The pairs plot shows protein intensity distribution of respective samples along the diagonal, with scatter plot and Pearson’s correlation coefficient of paired samples along the lower and upper off-diagonals respectively. The regression line in scatter plot is highlighted in red and significance of correlation coefficient is indicated with stars. **C)** The respective samples were grouped based on percentage distribution of principle component 1 on x-axis and principal component 2 on y-axis. **D)** The percentage co-efficient of variation (CV) distribution is plotted using a violin plot for BPA and mock group. The mean CV of two groups were indicated by red dotted line. **E)** The number of precursors, peptides, proteins and protein groups identified in individual samples is represented using log-scale on y-axis. **F)** The volcano plot displayed the comparison between BPA vs mock group with log2 fold change (FC) along x-axis and -log10 q-value along y-axis. The orange and dark-green circles show up and downregulate proteins respectively, with circle size corresponding to FC. The q-value cutoff of ≤0.05 and absolute log2 FC cutoff of ≥0.378 is highlighted with red dotted lines.

We then performed principal component analysis which distinctly separated the mock from BPA group with maximum PC1 variation of 20.94% on y-axis and PC2 variation of 16.79% on x-axis (**Figure 1C**). The median CV (coefficient of variation) distribution for mock and BPA group is 17.4% and 14.9% respectively (**Figure 1D**). Spectronaut generated de-novo library compiled 26,577 precursors mapping 22,471 peptides from 6273 unique protein ids extracted from 3592 protein groups. The individual detection of precursors, peptides, proteins and protein groups for each sample were listed in **Figure 1E**.

Next, we performed differential expression analysis of proteins between the BPA and Mock condition. To determine the fold change and FDR adjusted p-value (q-value) cutoff we plotted a distribution histogram and identified the upper and lower threshold of the distribution using inter-quartile range (IQR) calculation (**Figure S1A and S1B**). The calculated cutoff identified the proteins with log2 FC ≤-0.274 and ≥0.343 as differentially regulated. However, we used a classical FC limit of ≥1.3-fold (absolute Log2FC 0.378) and q-value threshold of ≤0.05 for identification of significantly regulated proteins between the groups (**File S1**). The comparison of BPA vs mock condition yielded 84 differentially expressed proteins (DEPs) among which 20 are down-regulated (dark green) and 64 are up-regulated (orange), as depicted in the volcano plot (**Figure 1F**). The Z-score abundance of the DEPs was plotted using a heat-map where orange and dark-green colours indicate higher and lower Z-score values respectively (**Figure 2A**). The heat-map separates two distinct clusters of proteins based on their regulation pattern.

**Figure 2:**
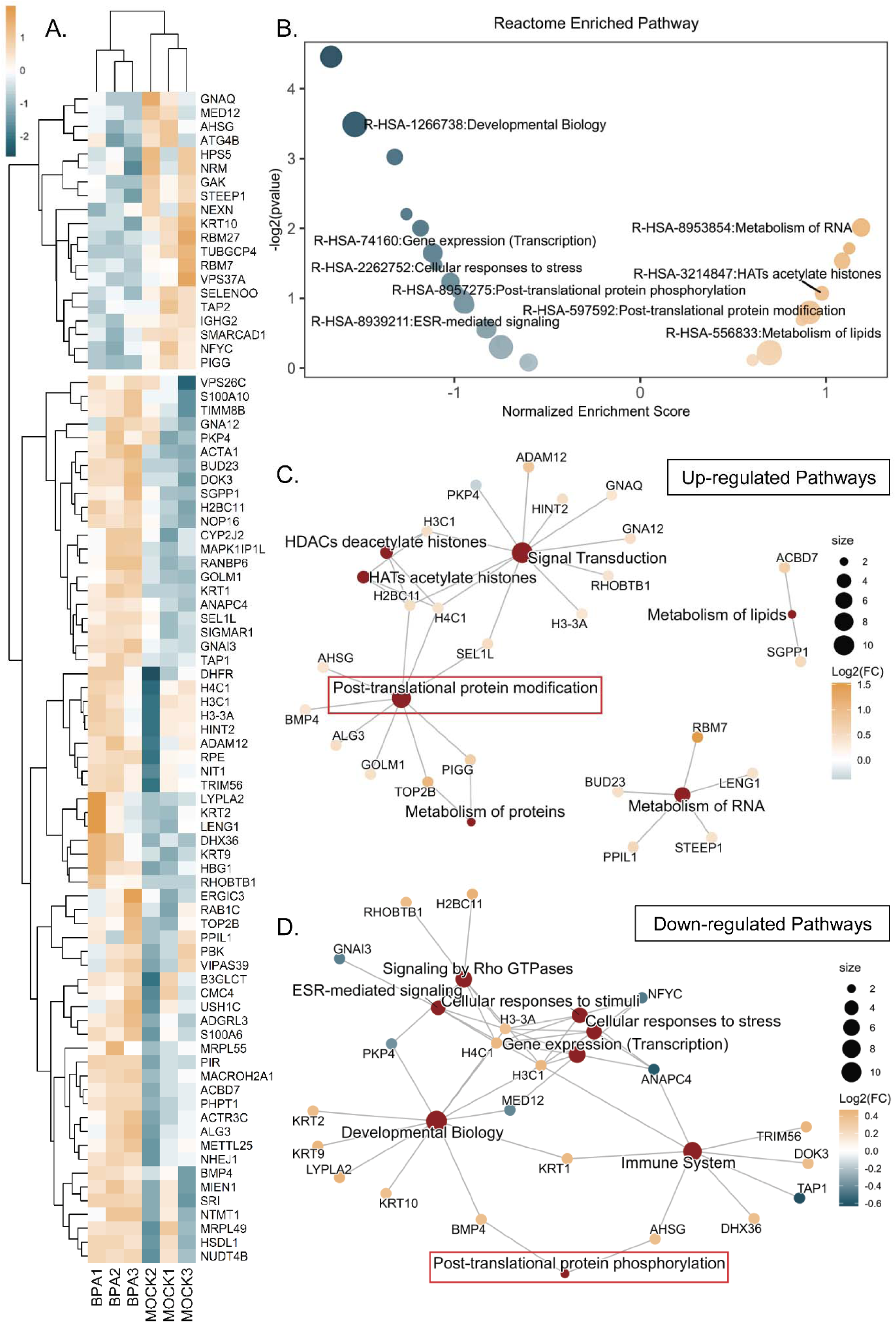
The dynamic alteration of global proteome post-BPA exposure. **A)** Heatmap revealed the differential protein expression pattern using z-score abundance where orange and dark-green color indicates higher and lower z-score values. **B)** The gene set enrichment analysis (GSEA) of differential proteins, enriched Reactome pathways represented using volcano plots. The normalized enrichments scores and -log2 p-value is plotted on x-axis and y-axis respectively. The same color scheme is used to represent the up and down regulated pathways. The circle size corresponds to number of proteins associated with the pathways. The category network (cnet) plot showed **C)** upregulated or **D)** downregulated Reactome pathways in dark-red nodes with edges connecting to their associated protein nodes coloured according to the log2 fold change. The node size of pathways corresponds to number of associated proteins. The relevant pathways are highlighted with red squares.

Further, to decipher the functional relevance of the regulated proteins we performed gene set enrichment analysis using Reactome database. The DEPs enriched 85 up-regulated and 18 down-regulated pathways (**File S1**) depicted in **Figure 2B**, where the colour dark-green and orange represented down-regulated and up-regulated pathways with circle size corresponding to number of genes involved in the pathway. Some highly relevant elevated pathways are Metabolism of RNA, HATs acetylate histones, post-translational protein modification and Metabolism of lipids. Whereas, Developmental Biology, Gene expression (Transcription), Cellular responses to stress, ESR-mediated signalling and post-translational protein phosphorylation are some down-regulated pathways that enriched upon BPA exposure. We next explored the potential biological complexity of BPA toxicity using category network (cnet) plot. The cluster of elevated pathways in cnet plot showed association of single proteins with multiple pathways, which are exclusively elevated in treatment group (**Figure 2C**). The down-regulated pathways also showed connection of single proteins with multiple pathways but, here the protein expression shows both up and down-regulatory trend (**Figure 2D**).

### 3.2. Bisphenol-A mediated trophoblast cell injury affects phosphoprotein-signatures

Cellular exposure to bisphenol-A often triggers specific phosphorylation events required for dynamic downstream signaling^32–36^. BPA administration induced dynamic proteome changes in trophoblast cells which deregulated pathways of post-translational protein modification and post-translational protein phosphorylation (**Figure 2B-D**). Thus, deciphering the BPA altered phospho-proteome signature become important for correlating the molecular mechanism of toxicity. We followed a metal affinity (TiO_2_) based enrichment procedure for phospho-peptide entrapment in tandem with information dependent data acquisition using ZenoTOF 7600 MS for phospho-peptide identification (**Figure 3A**). The peptide spectrum matches (PSMs) were scored in Fragpipe using hyper-score and next-score for confident phospho-site identification. The scatter plot distribution of both phospho-modified and nonmodified PSMs with its corresponding hyper-score in y-axis and next-score in x-axis shows distinct separation of phospho-modified PSMs along the diagonal in all the samples (**Figure 3B**). This confidently identified phospho PSMs were further filtered with 1% FDR cutoff and ultimately mapped onto a phospho-peptides. This information is further used for identification of phospho-site and phospho-proteins in BPA and mock condition separately.

**Figure 3:**
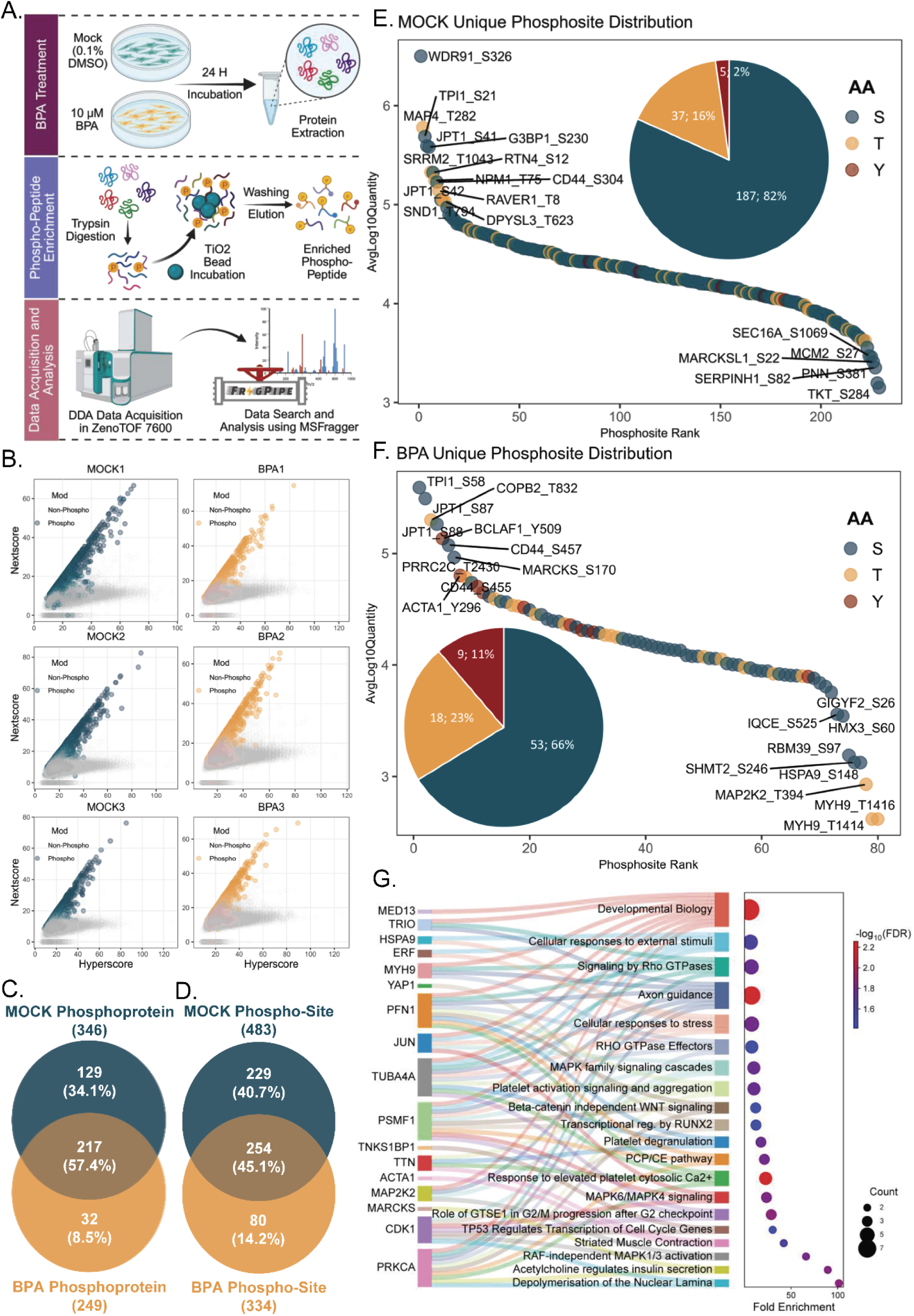
Reshaped phospho-proteome landscape due to bisphenol-A toxicity. **A)** The schematic pipeline followed for phospho-proteome enrichment and identification. **B)** The next-score and hyper-score for both phosphorylated and non-phosphorylated peptide spectrum matches (PSMs) were represented on y- and x-axis respectively with individual scatter plot for each sample. The Venn diagram displayed overlap between mock and BPA identified **C)** phospho-proteins and **D)** phospho-sites with numbers and percentage mentioned in brackets. The uniquely present phospho-sites in **E)** mock or **F)** BPA group were visualized with ranked abundance plot. The phospho-site rank in x-axis is plotted against its average log10 quantity on y-axis and the dots are coloured according to the phosphorylated residue. A pie chart inside the plot shows the distribution of phosphorylation on serine, threonine or tyrosine in mock or BPA condition. **G)** Shankey diagram of top 20 Reactome pathways enriched by BPA unique phospho-proteins. The dot size indicates protein counts while the adjusted p-value is represented using colors.

We identified a total 346 phospho-proteins in mock and 249 phospho-proteins in BPA condition, from which 217 (57.4%) phospho-proteins were common between the groups. We found 129 (34.1%) unique phospho-proteins in mock condition whereas in BPA there are only 32 (8.5%) unique phospho-proteins (**Figure 3C and File S2**). Subsequently, we reported a total of 483 and 334 phospho-sites in mock and BPA group respectively and found 254 (45.1%) common phospho-sites between them (**Figure 3D**). Finally, we extracted 229 (40.7%) unique phospho-sites in mock and 80 (14.2%) unique phospho-site in BPA samples, along with their respective average log10 quantity (**File S2**). We utilized this extracted matrix for plotting ranked abundance to describe the distribution of uniquely identified phospho-site in each comparison group. We identified mock-unique phosphorylated sites modified on serine (82%, 187), threonine (16%, 37), or tyrosine (2%, 5) are distributed with similar abundance (**Figure 3E**). However, in case of BPA we detected tyrosine modified phospho-sites (11%, 9) are more abundant than serine (66%, 58) or threonine (23%, 18) modified residues (**Figure 3F**).

Next, we performed pathway enrichment using over-representation analysis (ORA). The BPA unique phospho-proteins enriched pathways of Cellular response to external stimuli or stress, Signalling by Rho-GTPases, RHO GTPase Effectors, MAPK family signalling, MAPK6/MAPK4 signalling, Beta-catenin independent WNT signalling. The phospho-modified proteins primarily involved in these pathways include PRKCA, CDK1, MAP2K2, TTN, PSMF1, TUBA4A, JUN, PFN1, and MYH9 (**Figure 3G and File S2**). However, the unique phospho-proteins in mock condition represented the enriched pathways of cell cycle, vesicle mediated transport, membrane trafficking, mitotic metaphase and anaphase, golgi to ER and ER to golgi retrograde transport. These pathways implicated proteins including LMNB1, SPTAN1, NUP98, NUP153, GBF1, COPA, TUBA1B, ARFGAP2, NCBP1, SEC16A, ANAPC4, PDS5B, and CDC23, which were uniquely phosphorylated in the mock group (**Figure S1C and File S2**).

### 3.3. Global proteome enriched kinome map highlights BPA affected kinases

The maintenance of a protein phosphorylation signatures depends upon the intricate balance between kinase and phosphatase activity^52^. However, the phosphatase activity is more promiscuous with requirement of other regulatory sub-units, but kinases are precise often requiring specific sequences for substrate recognition^53^. Therefore, we took a more logical approach of connecting altered phospho-sites with their known target kinases. We started from exporting the list of known human kinases, which we then overlapped with our LC-MS/MS identifications to highlight the 75 kinases in our MS data (**Figure S2A and File S3**). These 75 kinases enriched multiple relevant pathways linked to BPA exposure, like ErbB signalling, EGFR tyrosine kinase inhibitor resistance, endocrine resistance, oxytocin signalling, MAPK signalling, Ras signalling and PI3K-Akt signalling (**Figure S2B and File S3**). Next, we explored the abundance distribution and regulation of these kinases in BPA vs mock comparison (**Figure 4A-B**). Interestingly, we observed that the kinases are equally distributed among the proteins from higher to lower abundance but does not show a distinct regulation between the comparison group. Therefore, we followed a stepwise selection method for identification of the regulated kinases (**Figure 4C**).

**Figure 4:**
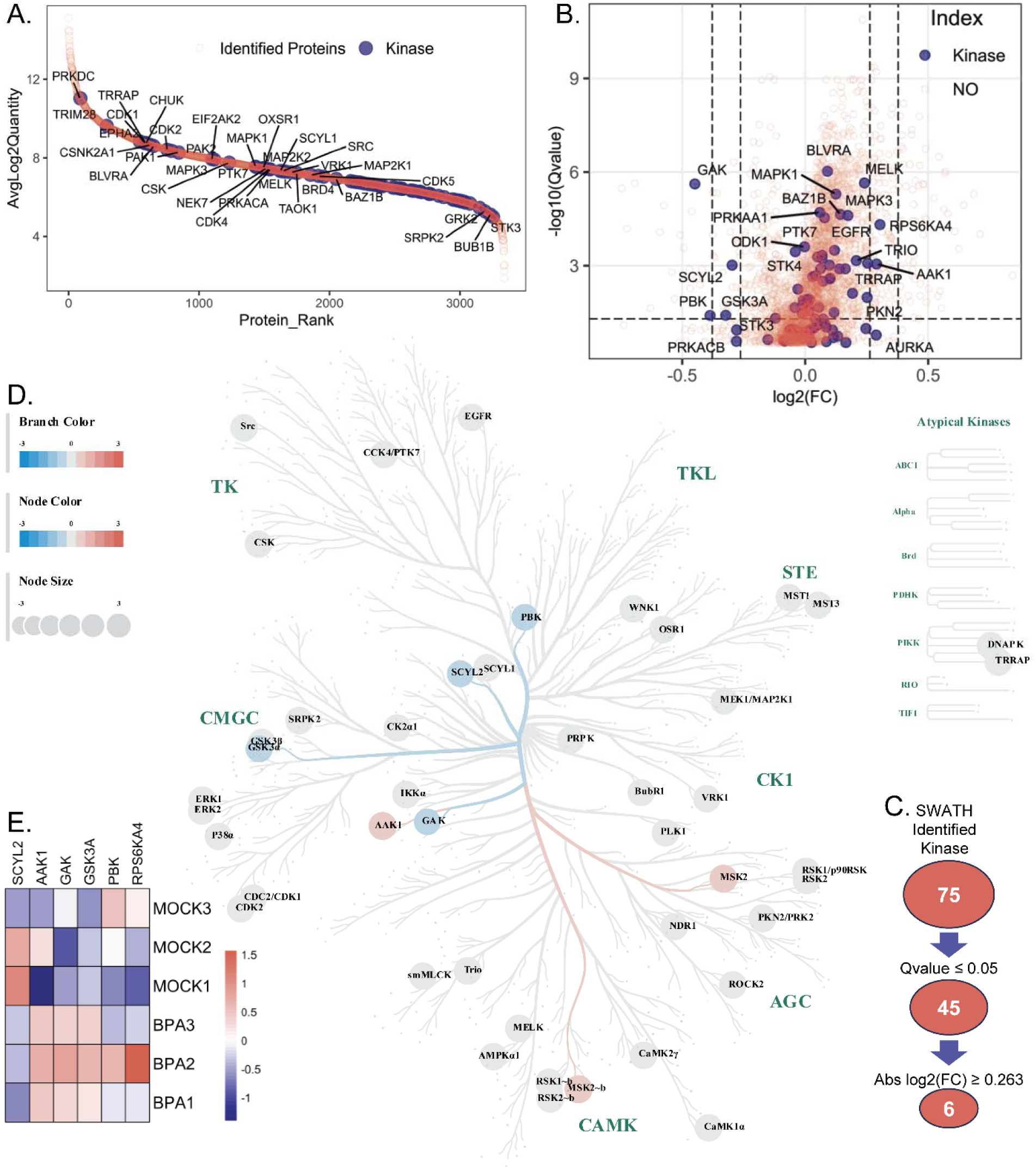
Early trimester trophoblast kinome homeostasis is affected by bisphenol-A: **A)** The ranked abundance plot highlights the global proteome identified kinase in blue. **B)** The volcano plot shows the differential information of the identified kinases with q-value filtering of ≤0.05 and absolute fold change cut-off of ≥1.2 and ≥1.3 highlighted with black dotted line. **C)** The filtering schema for identification of BPA regulated kinases. **D)** The q-value filtered kinases are visualized using kinome tree where kinase branches coloured with blue shows downregulation, while red branches indicate up-regulation. The gray nodes display identified kinases which are not regulated. **E)** The heatmap represents normalized expression values of the six regulated kinases in each BPA and MOCK samples with red color indicating higher expression and blue color indicating lower.

First, the q-value filtered kinases were visualized in human kinome tree using CORAL visualization tool to reveal the kinase distribution across all the known kinase families (**Figure 4D**). The coloured nodes highlighted the up-regulated (red) or down-regulated (blue) kinases that crossed the absolute fold change cutoff of ≤0.263, while other identified but not regulated kinases were marked with grey circles. The six significantly regulated kinases in BPA vs mock comparison were SCY1-like protein 2 (SCYL2), AP2-associated protein kinase 1 (AAK1), Cyclin-G-associated kinase (GAK), Glycogen synthase kinase-3 alpha (GSK3α), Lymphokine-activated killer T-cell-originated protein kinase (PBK) and Ribosomal protein S6 kinase alpha-4 (RPS6KA4/MSK2). The standardized abundance of regulated kinases was represented in a heatmap to reveal the over expression of AAK1, RPS6KA4 and the downregulation of GSK3α, GAK, PBK, SCYL2 in BPA treated condition (**Figure 4E**).

### 3.4. Bisphenol-A promotes kinase driven phospho-site regulation

The specificity of a kinase for a phosphorylation motif is usually conserved and depends upon the peptide recognition and its spatial availability^53^. Therefore, identification of target phospho-sites and its conserved motif for the regulated kinase become crucial. We extracted this information from PhosphoSitePlus for all the regulated kinases (**Figure 5A**). The kinase specific sequence motifs were enriched with conserved amino acids over the upstream (+7) and downstream (-7) flanking region around the central phosphorylation site, followed by identifications of known phospho-sites of these regulated kinases (**File S3**). GSK3α (186) enriched most known phospho-sites followed by PBK (22) and RPS6KA4 (MSK2) (12), while for GAK and AAK1 enriched only 6 and 4 known phospho-sites, respectively (**Figure S2C-G and File S3**). SCYL2 did not enriched any known phospho-site or motif sequence conserved for phosphorylation. We next combined all the extracted phospho-site information and removed the redundant signatures to accumulate 113 known sites. An upset plot was used to visualization the overlap of this known phospho-signatures with the uniquely identified phospho-sites in either mock or BPA group (**Figure 5B**). We found overlap of c-JUN phospho-site S63, which was uniquely present in BPA exposure, with the literature known phospho-signatures. This phosphorylation of c-JUN on serine 63 is reported by multiple kinases and PBK, which gets down-regulated upon BPA exposure, is one of them. Similarly, we reported the presence of tyrosine 279 phosphorylation on GSK3α uniquely in mock samples and when checked for known phospho-sites overlap, it revealed an auto-phosphorylation event of GSK3α on its tyrosine residue at 279.

**Figure 5:**
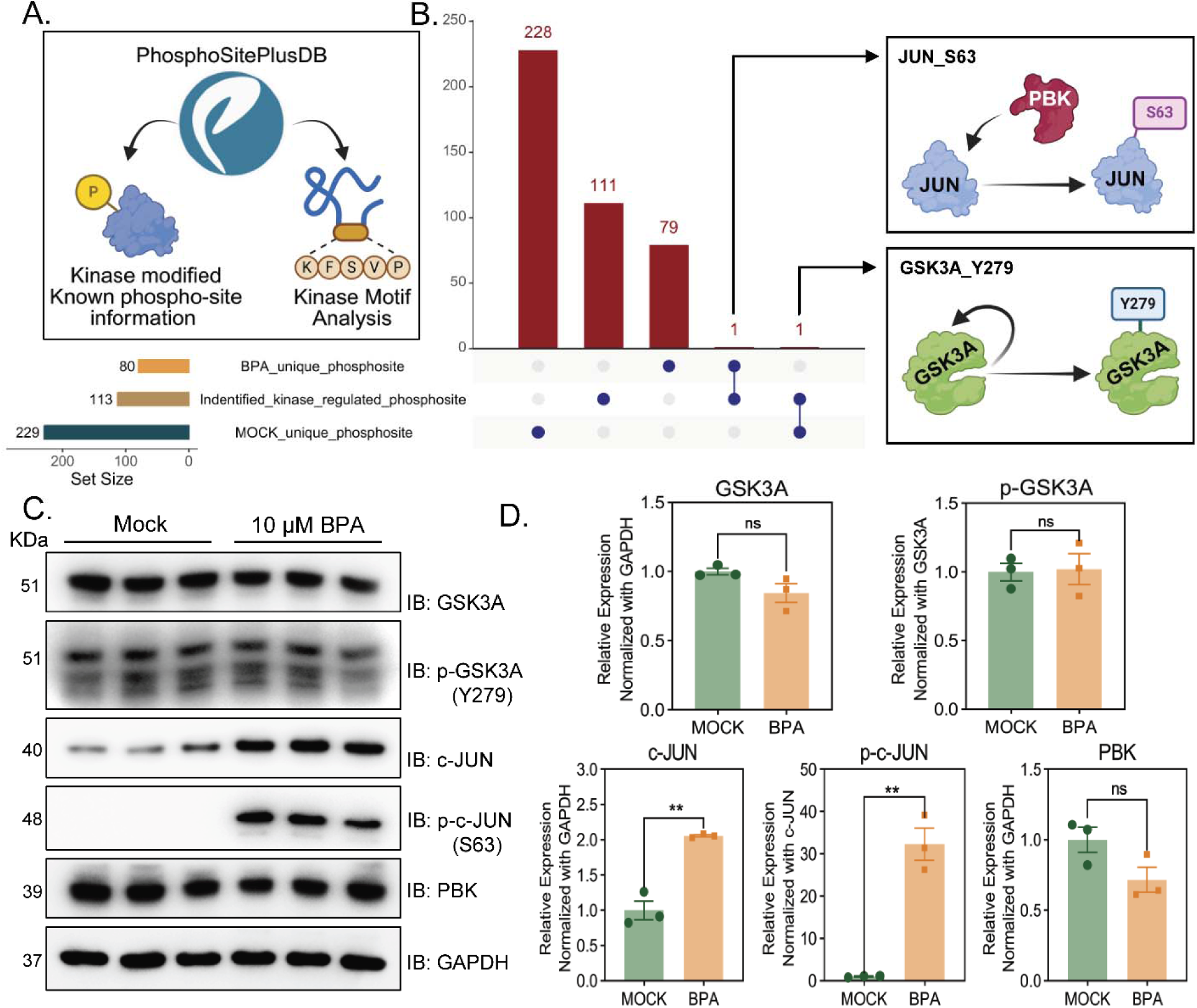
Identification of BPA altered phosphorylation pattern of c-JUN and GSK3A in HTR8/SVneo cells: **A)** schematic diagram displaying enrichment of regulated kinase identified motifs which modified the reported phospho-sites. **B)** The upset plot overlapped the kinase regulated known phospho-sites with either mock or BPA unique phospho-signatures. The intersection of phospho-c-JUN (S63) and phospho-GSK3A (Y279) along their target kinases are graphically represented. **C)** The western blot analysis is performed with three biological replicates of mock or BPA treated samples probed with GSK3α, phospho-GSK3α (Y279), c-JUN, phospho-c-JUN (S63), and PBK. GAPDH is used as loading control. **D)** The relative expression is normalized using intensity of either GAPDH or total non-phospho form and represented as mean ± SEM (N=3). The evaluation of statistical significance was performed by Student’s t-test. p-value of less than 0.05 is considered as significant.

These connections of regulated kinases with their corresponding unique phospho-signatures in either mock or BPA condition were further validated in HTR8/SVneo cells exposed to 10 μM BPA. We probed for phospho GSK3α (Y279) and phospho c-JUN (S63) along with their non-phospho forms using western blot and checked the kinase expression of PBK and GSK3α to find a holistic view of the regulatory kinetics (**Figure 5C**). This experiment revealed a non-significant down-regulation trend for GSK3α, but the phospho-Y-279-GSK3α expression remained unchanged. However, both phospho-S-63-c-JUN and total c-JUN abundance showed significant up-regulation upon BPA treatment. We also found a down-regulatory expression pattern for PBK similar to our proteomics data, but it also remained statistically non-significant (**Figure 5D**).

### 3.5. Dysregulation of GSK3α and c-JUN phosphorylation by BPA can cause placental toxicity

Pregnant women spontaneously exposed to BPA by ingestion, inhalation or dermal contact are at high risk of placental injury by BPA, as it can easily cross the placental barrier from maternal circulation. Placental heterogeneity plays a crucial role on how the BPA affect the placental homeostasis. Different lineages of trophoblast cells, like syncytiotrophoblasts and extravillous trophoblast, responds to chronic BPA dosage differently. Thus, it is important to validate the isolated cell culture findings in complex tissue system of heterogeneous cell types for establishing stronger evidence of toxicity mechanism. Therefore, we established a pregnancy mice model with BPA exposure by following a modified protocol from previously published study^30^. We orally administrated 200 μg/kg body-weight BPA every day in a mixture of ethanol and corn oil for 2 weeks before mating and after successful mating we continued the treatment until the dams were humanely euthanized at embryonic day 12.5 for placenta collection (**Figure 6A**). Mock control group were administered only with the mixture of ethanol and corn oil orally every day. Upon closer visual inspection of the uterine horn from both the mock and BPA exposed dams we didn’t observed any distinct morphological changes (**Figure 6B-C**). However, to validate the altered phosphoproteome signature revealed by the BPA treated HTR8/SVneo cellular model, we dissected the embryos to harvest only the placental tissue which were further used for western blot analysis. We took 2 placenta each from right and left part of the uterine horn for a single dam. We checked the phospho-Y-279 GSK3α, total GSK3α, phospho-S-63 c-JUN, total c-JUN and PBK expression in both BPA and mock treated placental samples (**Figure 6D**). Analysis of these placental blots revealed a significant downregulation of total GSK3A abundance and a significant up-regulation of total c-JUN expression. However, the phospho-forms, which is normalized using the total abundance of the respective non-phospho proteins, showed no significant regulation. The same observation is reported for PBK expression also (**Figure 6E**).

**Figure 6:**
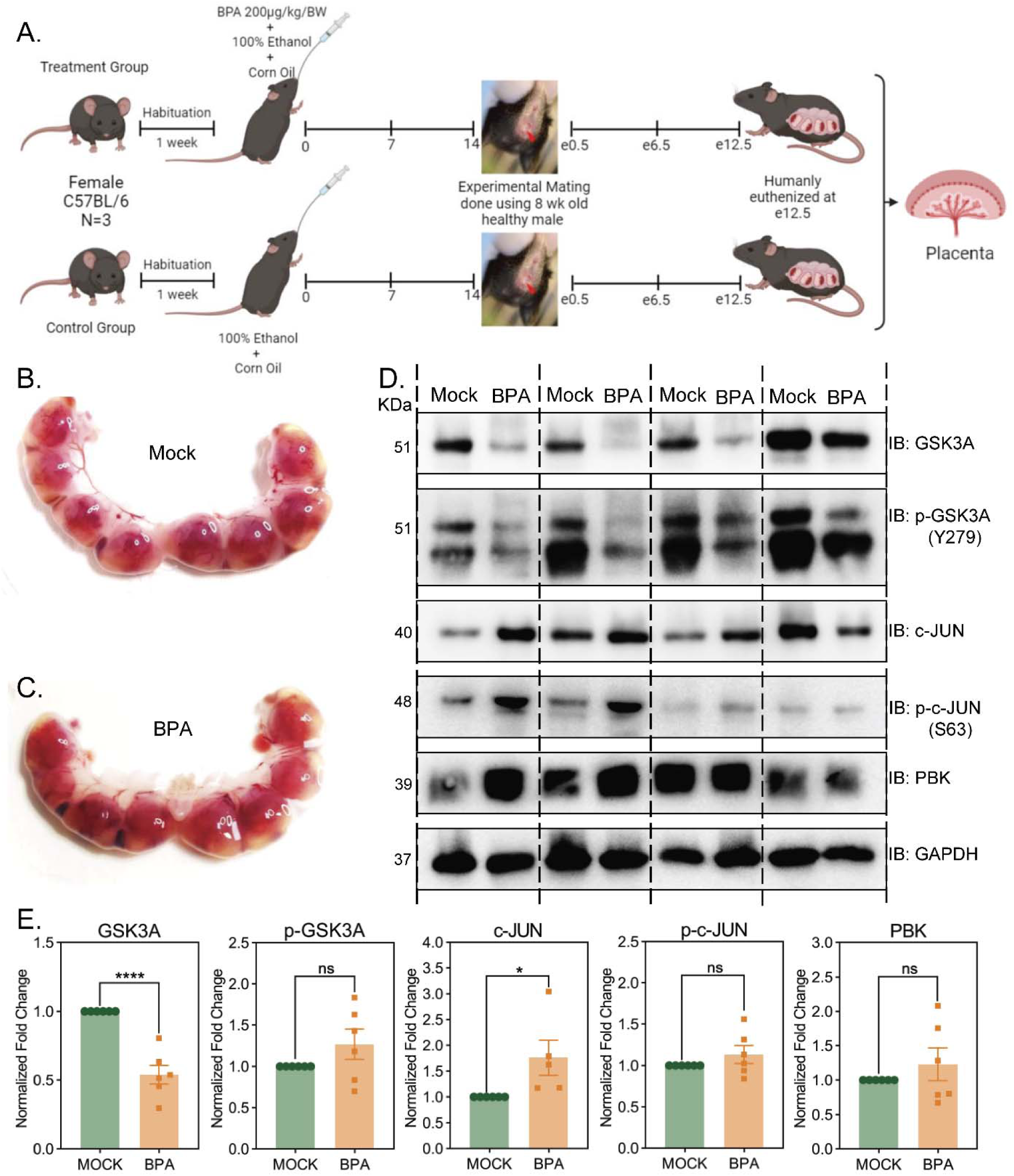
The BPA altered phospho-signatures in trophoblast cells are validated in BPA administered mice placenta samples. **A)** The workflow followed for generating BPA exposed pregnancy model in mice. The uterine horn showing the embryos with fetus and placenta inside were collected from either **B)** mock or **C)** BPA exposed mice post euthanization and then imaged. **D)** The western blot analysis of GSK3α, phospho-GSK3α (Y279), c-JUN, phospho-c-JUN (S63), and PBK is performed using placentas (N=6) from mock and BPA exposed mice. Here also, equal loading is indicated using GAPDH. **E)** The densitometric analysis of the blots are displayed as mean ± SEM (N=6). The p-value of <0.05, calculated using Student’s t-test, is considered statistically significant.

## 4. DISCUSSION

Environmental exposure of toxic pollutants is one of the major health concerns for mothers worldwide^8^. Pollutants, such as endocrine disrupting chemicals (EDCs) can affect both maternal and fetal health during the pregnancy due to their constant presence in daily use items and potent impact over biological functions^9^. Bisphenol-A (BPA), a plastic or epoxy resin leachate, act as an endocrine disrupting chemical by triggering hormonal imbalance during gestation and causing fetal injury. Elevated presence of BPA in maternal circulation was associated with pregnancy related disorders like pre-eclampsia^14–16^ and preterm birth^11,12^. This BPA in maternal circulation can readily cross the placental barrier and impart cellular damage to the trophoblast cells in placenta. Although, multiple studies investigated the mechanism of BPA toxicity in placenta using cellular model of placental lineage like BeWo, JEG-3 or HTR8/SVneo, but how early BPA exposure can harm the placental axis is still less explored. Thus, we exposed the first trimester extra-villous trophoblast cells with physiologically relevant concentration of BPA and evaluated its proteome wide differential pattern to reveal the BPA specific alteration of cellular signalling. We employed a SWATH based proteomics workflow to acquire data, that demonstrates a high pairwise Pearson’s correlation with biologically relevant CV distribution and distinct separation of BPA and mock group in principle component analysis (**Figure 1A-D**). The ZenoSWATH based quantitation of the identified protein groups (3335) across the samples displayed a distinct regulation of 84 proteins with specified cutoff in BPA vs mock comparison (**Figure 1F and File S1**). This differentially expressed proteins were distributed perfectly in two separate clusters of up (64) or down (20) regulated proteins (**Figure 2A**) showing a dynamic pattern of proteome change in early trimester trophoblast cells post-BPA exposure. Pathway enrichment captured down-regulation of BPA affected pathways reported in previous studies, such as epigenetic regulation of gene expression^19^, signalling by WNT^19,29^, oxidative stress induced senescence^18,24^, signalling by nuclear receptors^54^, ESR-mediated signaling^13,21^, and gene expression (Transcription)^25^. The downregulation of pathways for developmental biology and reproduction provide important insight relevant to BPA exposure. Similarly, literature reported up-regulated pathways includes metabolism of lipids^55^, signalling by NOTCH^56^, HATs acetylate histones^57^ and post-translational protein modification^57^ (**Figure 2B and File S1**). Interestingly, pathway enrichment revealed that, BPA exposure down-regulates post-translational protein phosphorylation in first trimester trophoblast cells (**Figure 2D**). This observation directed us to explore phospho-proteome dynamics in BPA exposed trophoblast cellular model using TiO_2_-based phospho-peptide enrichment in tandem with high throughput LC-MS/MS analysis.

The phospho-proteome experiment enriched a smaller number of unique phospho-site (80) and phospho-proteins (32) in BPA treated condition, than mock enriched unique phospho-site (229) and phospho-proteins (346) (**Figure 3C-D and File S2**), in line with our previous pathway enrichment analysis. However, unique phospho-site from both BPA and mock groups were majorly modified on serine residue followed by modification on threonine and tyrosine residue (**Figure 3E-F**). Next, we observed that multiple pathways enriched by BPA unique phospho-proteins overlapped with differential proteome enriched pathways, such as developmental biology, cellular responses to stress, signalling by Rho GTPases, RHO GTPase effectors and transcriptional regulation by RUNX1 (**Figure 3G and File S2**). BPA exposure uniquely affected phospho-proteins involved in MAPK family signalling cascades, MAPK6/MAPK4 signalling, and RAF-independent MAPK1/3 activation pathways substantiating the role of altered MAPK family protein phosphorylation in BPA dependent regulation of trophoblast toxicity^58^. The similar pathway enrichment with mock unique phospho-proteins enriched the pathways required for healthy survival of cells, like vesicle-mediated transport, cell cycle, membrane trafficking, Golgi-to-ER and ER-to-Golgi transport, SUMOylation of RNA binding proteins and transport of mature transcript to cytoplasm (**Figure S1C**). This phospho-sites were carefully regulated by activity of kinases and phosphatase^52^. However, the kinase activity is more substrate and motif specific^53^ which directed us to isolate 75 kinases from the SWATH identified global proteome data (**Figure 4A-B and File S3**). These kinases were further filtered using q-value and log2 fold change cutoff, which identified six kinases significantly regulated upon BPA exposure (**Figure 4C-D**). The up-regulated kinases are AAK1 and RPS6KA4 from NAK and AGC kinase family respectively. Whereas the GSK3α (CMGC kinase family), PBK (TOPK subfamily), SCYL2 (SCY1 subfamily) and GAK (NAK subfamily) were found down-regulated (**Figure 4E**). These regulated kinases have selection towards specific motifs and targets, which we extracted from public databases. The AAK1 and GAK shows specificity only for phosphorylation at threonine reside (**Figure S2E-F**), whereas RPS6KA4 (MSK2) phosphorylates serine residues only (**Figure S2G**). Contrary to this, PBK and GSK3α phosphorylates both residues but shows more affinity towards serine (**Figure S2C-D**). This extracted known phospho-sites for BPA regulated kinases **(File S3**) were finally overlapped with unique phospho-sites identified in mock or BPA samples. We observed only phospho-c-JUN (S63) from BPA and phospho-GSK3α (Y279) from mock samples are common with kinase regulated known phospho-sites (**Figure 5B**). Further investigation for these phospho-site associated kinases identified PBK which phosphorylates c-JUN at serine 63^59^, and GSK3α which auto-phosphorylates itself at 279 position tyrosine residue^60^. Both the phosphorylation event act as a “on” switch required for activation of respective protein function. The auto-phosphorylation of GSK3α at Y279 is essential for its stability and increased catalytic activity^61^. However, c-JUN phosphorylation at serine 63 drives transcriptional activation of AP1 complex elevating c-JUN target gene expression^62^. Therefore, we validated the dynamic regulation of this kinase mediated phospho-protein expression in both BPA administered cellular and murine model.

We observed no significant changes of GSK3α, phospho-GSK3α (Y279) and PBK expression in HTR8/SVneo cells post BPA exposure. However, c-JUN and phospho-c-JUN (S63) expression were significantly up-regulated by 2-fold and 30-fold respectively (**Figure 5C-D**). We even detected the phospho-c-JUN (S63) bands exclusively in BPA exposed samples, which suggest that PBK expression is not the only factor in this complex signalling cascade. Either the altered activity of PBK, or other kinases responsible for c-JUN S63 phosphorylation is playing a major role in the background. The placentas from BPA administrated pregnant mice, although did not show any morphological differences when compared with mock placentas, but revealed altered phospho dynamics at molecular level (**Figure 6B-D**). We found significant protein level down-regulation of total GSK3α and upregulation of total c-JUN expression. The expression of PBK and phospho-forms of GSK3α and c-JUN does not show any statistical differences (**Figure 6E**). This potentially indicates that the phosphorylation of c-JUN (S63) and GSK3α (Y279) is increased due to over-expression of their total protein abundance and not due to altered kinase activity. However, to determine the downstream effect of c-JUN phosphorylation we extracted target gene information for both mice and human c-JUN protein (**Figure S3A and C**), followed by their pathway enrichment. which highlighted conserved pathways like activation of matrix metalloproteinases, extracellular matrix (ECM) organization, degradation of extracellular matrix and collagen degradation in both genus (**Figure S3B and D; File S4**). These are essential functions disrupted by BPA exposure during early stages of placentation.^17^ Whereas, GSK3α is important for cellular metabolism, survival and proliferation^63^ which can be dysregulated by bisphenol-A^64^ during gestation, imparting chronic placental injury. Therefore, our study suggest that BPA can cause detrimental effect on first trimester trophoblast cells during early placentation events through altered phosphorylation dynamics of c-JUN and GSK3α. This can affect the downstream signalling for ECM remodelling, proliferation and survival, essential for maintenance of placental homeostasis and healthy fetal growth. However, this study does not explain how BPA is altering c-JUN and GSK3α phosphorylation and further limits its observation by only exploring cellular and murine models. In future, correlating this finding with adverse pregnancy outcomes will add more insights, highlighting a mechanistic model for BPA mediated placental toxicity.

## 5. ENVIROMENTAL IMPLICATIONS

Rapid industrial and urban growth over last few centuries have polluted our environment with various toxic chemicals. Daily exposure to these pollutants from everyday house-hold items have caused a serious distress on our health. Endocrine disrupting chemicals (EDCs) are one such toxic pollutants which interferes with our healthy hormonal homeostasis. This made EDCs an alarming concern for pregnant mothers due of their susceptibility towards hormonal imbalance mediated through EDCs. One such endocrine disrupting chemical is bisphenol-A (BPA) which mimics estrogen structure and can impart damage on the developing feto-maternal unit via alteration of hormonal signalling. The potent and persistent environmental exposure of BPA from plastic goods can cause BPA accumulation in maternal circulation through ingestion, inhalation or dermal contact. During gestation, this BPA can cross the placental barrier and can induce detrimental effect on the growing fetus. This BPA mediated feto-placental injuries can provoke adverse pregnancy outcomes leading to neonatal complications. However, the molecular mechanism of BPA toxicity on feto-placental axis during pregnancy needs more study. Thus, we mapped the BPA altered regulation of global proteome and phospho-proteome signatures in early trimester trophoblast cells which highlighted dysregulation of kinase expression and function as a key mediator of BPA toxicity. The result suggested, an altered expression of phospho-GSK3α and phospho-c-JUN can affect downstream signalling in BPA exposed trophoblast or murine model. However, these results do not explain, if it is a cause or effect of BPA exposure and moreover, the findings are not validated using exposure studies in human. Even with these limitations, we are able to establish an altered phosphorylation mediated mechanism of BPA toxicity in placenta which may trigger adverse pregnancy outcome related complications during pregnancy.

## ASSOCIATED CONTENT

### Supplementary information

Supporting Information is available free of charge at http://pubs.acs.org

**Supplementary Figure S1:** Distribution of differential parameters and mock phospho-protein enriched pathways.

**Supplementary Figure S2:** Exploring the identified kinases.

**Supplementary Figure S3:** c-JUN target associated pathways.

**Supplementary File S1:** The differentially expressed proteins and their enriched Reactome pathways post BPA exposure in HTR8/SVneo cells.

**Supplementary File S2:** The mock and BPA unique phospho-proteins and their respective Reactome enriched pathways.

**Supplementary File S3:** The 75 SWATH identified kinases with their KEGG pathway enrichment. The regulated kinases were highlighted in yellow with their known targets of phosphorylation.

**Supplementary File S4:** The Reactome pathway enrichment of human and mice c-JUN target proteins.

### Data Availability

The mass spectrometry data are available online through the ProteomeXchange Consortium via the PRIDE (https://www.ebi.ac.uk/pride/) partner repository with the data set identifier PXD074780 (**Reviewer access details: Project accession:** PXD074780; **Token:** dgiY1vcl625y)

## AUTHOR INFORMATION

### Author Contributions

TKM and AB conceived the project and designed the experiments. AB, NK and SS performed phospho-proteomics and proteomics sample preparation and data acquisition. AB carried out bioinformatic analysis and wet-lab experiments. AB and TKM wrote the manuscript. All authors read and approve the final version of the manuscript.

### Notes

The authors declare no competing interest

## ACKNOWLEDGMENTS

TKM acknowledges the Regional Centre for Biotechnology for its intramural research funding. We thank Dr. Pallavi Kshetrapal for providing us the HTR8/SVneo cell line. We express our sincere gratitude to all the technical staff at RCB for keeping the equipment in functional condition. We are grateful to all the members of Laboratory of Functional Proteomics for maintaining an excellent research environment. Sandhini Saha thanked ICMR, Ankit Biswas thanked DBT and NK thanks CSIR for fellowship.

---

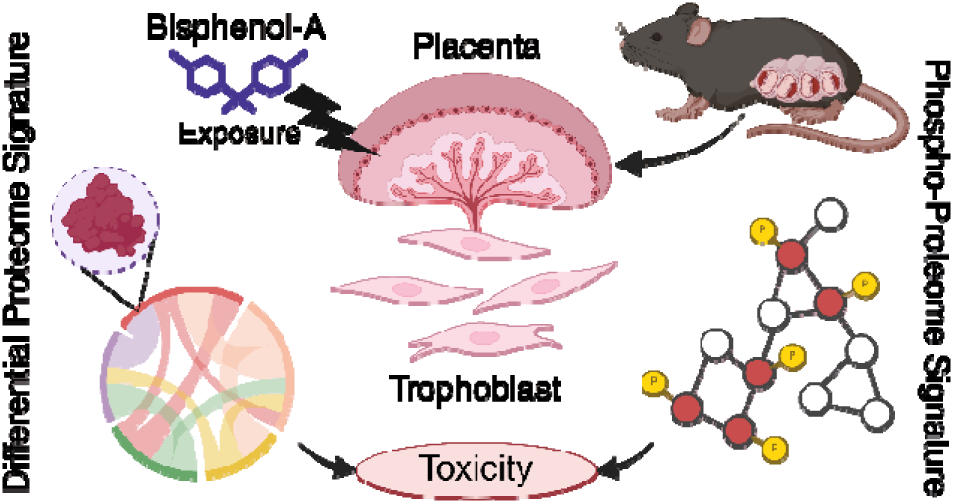
For Table of Contents Only.

## REFERENCES

(1) Burton, G. J.; Fowden, A. L. The placenta: a multifaceted, transient organ. Philosophical Transactions of the Royal Society B: Biological Sciences 2015, 370, 20140066.

(2) Howell, K. R.; Powell, T. L. Effects of maternal obesity on placental function and fetal development. Reproduction 2017, 153, R97–R108.

(3) Sferruzzi-Perri, A. N.; Camm, E. J. The programming power of the placenta. Frontiers in physiology 2016, 7, 33.

(4) Dimasuay, K. G.; Boeuf, P.; Powell, T. L.; Jansson, T. Placental responses to changes in the maternal environment determine fetal growth. Frontiers in physiology 2016, 7, 12.

(5) Mathiesen, L.; Buerki-Thurnherr, T.; Pastuschek, J.; Aengenheister, L.; Knudsen, L. E. Fetal exposure to environmental chemicals; insights from placental perfusion studies. Placenta 2021, 106, 58–66.

(6) Padula, A. M.; Monk, C.; Brennan, P. A.; Borders, A.; Barrett, E. S.; McEvoy, C. T.; Foss, S.; Desai, P.; Alshawabkeh, A.; Wurth, R.; others A review of maternal prenatal exposures to environmental chemicals and psychosocial stressors—implications for research on perinatal outcomes in the ECHO program. Journal of Perinatology 2020, 40, 10–24.

(7) Merrill, A. K.; Sobolewski, M.; Susiarjo, M. Exposure to endocrine disrupting chemicals impacts immunological and metabolic status of women during pregnancy. Molecular and cellular endocrinology 2023, 577, 112031.

(8) Hilz, E. N.; Gore, A. C. Endocrine-disrupting chemicals: science and policy. Policy insights from the behavioral and brain sciences 2023, 10, 142–150.

(9) Kawa, I. A.; Fatima, Q.; Mir, S. A.; Jeelani, H.; Manzoor, S.; Rashid, F.; others Endocrine disrupting chemical Bisphenol A and its potential effects on female health. Diabetes & Metabolic Syndrome: Clinical Research & Reviews 2021, 15, 803–811.

(10) Li, J.; Wu, C.; Zhao, H.; Zhou, Y.; Cao, G.; Yang, Z.; Hong, Y.; Xu, S.; Xia, W.; Cai, Z. Exposure assessment of bisphenols in Chinese women during pregnancy: a longitudinal study. Environmental science & technology 2019, 53, 7812–7820.

(11) Huang, S.; Li, J.; Xu, S.; Zhao, H.; Li, Y.; Zhou, Y.; Fang, J.; Liao, J.; Cai, Z.; Xia, W. Bisphenol A and bisphenol S exposures during pregnancy and gestational age– A longitudinal study in China. Chemosphere 2019, 237, 124426.

(12) Namat, A.; Xia, W.; Xiong, C.; Xu, S.; Wu, C.; Wang, A.; Li, Y.; Wu, Y.; Li, J. Association of BPA exposure during pregnancy with risk of preterm birth and changes in gestational age: A meta-analysis and systematic review. Ecotoxicology and Environmental Safety 2021, 220, 112400.

(13) Cao, Y.; Chen, Z.; Zhang, M.; Shi, L.; Qin, S.; Lv, D.; Li, D.; Ma, L.; Zhang, Y. Maternal exposure to bisphenol A induces fetal growth restriction via upregulating the expression of estrogen receptors. Chemosphere 2022, 287, 132244.

(14) Ye, Y.; Zhou, Q.; Feng, L.; Wu, J.; Xiong, Y.; Li, X. Maternal serum bisphenol A levels and risk of pre-eclampsia: a nested case–control study. The European Journal of Public Health 2017, 27, 1102–1107.

(15) Dagdeviren, G.; Arslan, B.; Keles, A.; Yücel Çelik, O.; Arat, Ӧ.; Caglar, A. T. Thë evaluation of serum bisphenol A in patients with preeclampsia. Journal of Obstetrics and Gynaecology Research 2023, 49, 1322–1327.

(16) Cantonwine, D. E.; Meeker, J. D.; Ferguson, K. K.; Mukherjee, B.; Hauser, R.; McElrath, T. F. Urinary concentrations of bisphenol A and phthalate metabolites measured during pregnancy and risk of preeclampsia. Environmental health perspectives 2016, 124, 1651.

(17) Müller, J. E.; Meyer, N.; Santamaria, C. G.; Schumacher, A.; Luque, E. H.; Zenclussen, M. L.; Rodriguez, H. A.; Zenclussen, A. C. Bisphenol A exposure during early pregnancy impairs uterine spiral artery remodeling and provokes intrauterine growth restriction in mice. Scientific reports 2018, 8, 9196.

(18) Sun, Y.; Sha, M.; Qin, Y.; Xiao, J.; Li, W.; Li, S.; Chen, S. Bisphenol A induces placental ferroptosis and fetal growth restriction via the YAP/TAZ-ferritinophagy axis. Free Radical Biology and Medicine 2024, 213, 524–540.

(19) Ye, Y.; Tang, Y.; Xiong, Y.; Feng, L.; Li, X. Bisphenol A exposure alters placentation and causes preeclampsia-like features in pregnant mice involved in reprogramming of DNA methylation of WNT2. The FASEB Journal 2018, 33, 2732.

(20) Wang, Z.; An, R.; Zhang, L.; Li, X.; Zhang, C. Exposure to bisphenol A jeopardizes decidualization and consequently triggers preeclampsia by up-regulating CYP1B1. Journal of Hazardous Materials 2025, 486, 137032.

(21) Adu-Gyamfi, E. A.; Rosenfeld, C. S.; Tuteja, G. The impact of bisphenol A on the placenta. Biology of Reproduction 2022, 106, 826–834.

(22) Balakrishnan, B.; Henare, K.; Thorstensen, E. B.; Ponnampalam, A. P.; Mitchell, M. D. Transfer of bisphenol A across the human placenta. American journal of obstetrics and gynecology 2010, 202, 393–e1.

(23) Guo, J.; Liu, K.; Yang, J.; Su, Y. Prenatal exposure to bisphenol A and neonatal health outcomes: A systematic review. Environmental Pollution 2023, 335, 122295.

(24) Ponniah, M.; Billett, E. E.; De Girolamo, L. A. Bisphenol A increases BeWo trophoblast survival in stress-induced paradigms through regulation of oxidative stress and apoptosis. Chemical Research in Toxicology 2015, 28, 1693–1703.

(25) Rajakumar, C.; Guan, H.; Langlois, D.; Cernea, M.; Yang, K. Bisphenol A disrupts gene expression in human placental trophoblast cells. Reproductive toxicology 2015, 53, 39–44.

(26) Li, X.; Wang, Y.; Wei, P.; Shi, D.; Wen, S.; Wu, F.; Liu, L.; Ye, N.; Zhou, H. Bisphenol A affects trophoblast invasion by inhibiting CXCL8 expression in decidual stromal cells. Molecular and Cellular Endocrinology 2018, 470, 38–47.

(27) Wei, P.; Ru, D.; Li, X.; Shi, D.; Zhang, M.; Xu, Q.; Zhou, H.; Wen, S. Exposure to environmental bisphenol A inhibits HTR-8/SVneo cell migration and invasion. Journal of Biomedical Research 2020, 34, 369.

(28) Yue, H.; Zhu, H.; Wu, X.; Tian, Y.; Zhang, J.; Hu, Y.; Ji, X.; Sang, N. Maternal bisphenol A (BPA) exposure induces placental dysfunction and health risk in adult female offspring: Insights from a mouse model. Science of the Total Environment 2025, 958, 177714.

(29) Tait, S.; Tassinari, R.; Maranghi, F.; Mantovani, A. Bisphenol A affects placental layers morphology and angiogenesis during early pregnancy phase in mice. Journal of Applied Toxicology 2015, 35, 1278–1291.

(30) Mao, J.; Jain, A.; Denslow, N. D.; Nouri, M.-Z.; Chen, S.; Wang, T.; Zhu, N.; Koh, J.; Sarma, S. J.; Sumner, B. W.; others Bisphenol A and bisphenol S disruptions of the mouse placenta and potential effects on the placenta–brain axis. Proceedings of the National Academy of Sciences 2020, 117, 4642–4652.

(31) Zha, X.; Elsabagh, M.; Zheng, Y.; Zhang, B.; Wang, H.; Bai, Y.; Zhao, J.; Wang, M.; Zhang, H. Impact of Bisphenol A exposure on maternal gut microbial homeostasis, placental function, and fetal development during pregnancy. Reproductive Toxicology 2024, 129, 108677.

(32) Wan, X.; Ru, Y.; Chu, C.; Ni, Z.; Zhou, Y.; Wang, S.; Zhou, Z.; Zhang, Y. Bisphenol A accelerates capacitation-associated protein tyrosine phosphorylation of rat sperm by activating protein kinase A. Acta Biochimica et Biophysica Sinica 2016, 48, 573–580.

(33) Li, N.; Kang, H.; Peng, Z.; Wang, H.-f.; Weng, S.-q.; Zeng, X.-h. Physiologically detectable bisphenol A impairs human sperm functions by reducing protein-tyrosine phosphorylation. Ecotoxicology and Environmental Safety 2021, 221, 112418.

(34) Fang, F.; Chen, D.; Yu, P.; Qian, W.; Zhou, J.; Liu, J.; Gao, R.; Wang, J.; Xiao, H. Effects of Bisphenol A on glucose homeostasis and brain insulin signaling pathways in male mice. General and comparative endocrinology 2015, 212, 44–50.

(35) Lee, C.-T.; Hsieh, C.-F.; Wang, J.-Y. Bisphenol a induces autophagy defects and AIF dependent apoptosis via HO-1 and AMPK to degenerate N2a neurons. International Journal of Molecular Sciences 2021, 22, 10948.

(36) Seki, S.; Aoki, M.; Hosokawa, T.; Saito, T.; Masuma, R.; Komori, M.; Kurasaki, M. Bisphenol-A suppresses neurite extension due to inhibition of phosphorylation of mitogen-activated protein kinase in PC12 cells. Chemico-biological interactions 2011, 194, 23–30.

(37) Saha, S.; Verma, R.; Kumar, C.; Kumar, B.; Dey, A. K.; Surjit, M.; Mylavarapu, S. V.; Maiti, T. K. Proteomic analysis reveals USP7 as a novel regulator of palmitic acidinduced hepatocellular carcinoma cell death. Cell Death & Disease 2022, 13, 563.

(38) Chambers, M. C.; Maclean, B.; Burke, R.; Amodei, D.; Ruderman, D. L.; Neumann, S.; Gatto, L.; Fischer, B.; Pratt, B.; Egertson, J.; others A cross-platform toolkit for mass spectrometry and proteomics. Nature biotechnology 2012, 30, 918–920.

(39) Kong, A. T.; Leprevost, F. V.; Avtonomov, D. M.; Mellacheruvu, D.; Nesvizhskii, A. I. MSFragger: ultrafast and comprehensive peptide identification in mass spectrometry–based proteomics. Nature methods 2017, 14, 513–520.

(40) Yang, K. L.; Yu, F.; Teo, G. C.; Li, K.; Demichev, V.; Ralser, M.; Nesvizhskii, A. I. MSBooster: improving peptide identification rates using deep learning-based features. Nature Communications 2023, 14, 4539.

(41) da Veiga Leprevost, F.; Haynes, S. E.; Avtonomov, D. M.; Chang, H.-Y.; Shanmugam, A. K.; Mellacheruvu, D.; Kong, A. T.; Nesvizhskii, A. I. Philosopher: a versatile toolkit for shotgun proteomics data analysis. Nature methods 2020, 17, 869–870.

(42) Yu, F.; Haynes, S. E.; Nesvizhskii, A. I. IonQuant enables accurate and sensitive labelfree quantification with FDR-controlled match-between-runs. Molecular & Cellular Proteomics 2021, 20, 100077.

(43) Ge, S. X.; Jung, D.; Yao, R. ShinyGO: a graphical gene-set enrichment tool for animals and plants. Bioinformatics 2020, 36, 2628–2629.

(44) Tang, D.; Chen, M.; Huang, X.; Zhang, G.; Zeng, L.; Zhang, G.; Wu, S.; Wang, Y. SRplot: A free online platform for data visualization and graphing. PloS one 2023, 18, e0294236.

(45) Vandenberg, L. N.; Chahoud, I.; Padmanabhan, V.; Paumgartten, F. J.; Schoenfelder, G. Biomonitoring studies should be used by regulatory agencies to assess human exposure levels and safety of bisphenol A. Environmental health perspectives 2010, 118, 1051.

(46) Dolinoy, D. C.; Huang, D.; Jirtle, R. L. Maternal nutrient supplementation counteracts bisphenol A-induced DNA hypomethylation in early development. Proceedings of the National Academy of Sciences 2007, 104, 13056–13061.

(47) Eid, S.; Turk, S.; Volkamer, A.; Rippmann, F.; Fulle, S. KinMap: a web-based tool for interactive navigation through human kinome data. BMC bioinformatics 2017, 18, 16.

(48) Metz, K. S.; Deoudes, E. M.; Berginski, M. E.; Jimenez-Ruiz, I.; Aksoy, B. A.; Hammerbacher, J.; Gomez, S. M.; Phanstiel, D. H. Coral: clear and customizable visualization of human kinome data. Cell systems 2018, 7, 347–350.

(49) Hornbeck, P. V.; Kornhauser, J. M.; Latham, V.; Murray, B.; Nandhikonda, V.; Nord, A.; Skrzypek, E.; Wheeler, T.; Zhang, B.; Gnad, F. 15 years of PhosphoSitePlusfi: integrating post-translationally modified sites, disease variants and isoforms. Nucleic acids research 2019, 47, D433–D441.

(50) Liska, O.; Bohár, B.; Hidas, A.; Korcsmáros, T.; Papp, B.; Fazekas, D.; Ari, E. TFLink: an integrated gateway to access transcription factor–target gene interactions for multiple species. Database 2022, 2022, baac083.

(51) Morice, L.; Benâıtreau, D.; Dieudonńe, M.-N.; Morvan, C.; Serazin, V.; de Mazancourt, P.; Pecquery, R.; Dos Santos, E. Antiproliferative and proapoptotic effects of bisphenol A on human trophoblastic JEG-3 cells. Reproductive toxicology 2011, 32, 69–76.

(52) Nasa, I.; Kettenbach, A. N. Coordination of protein kinase and phosphoprotein phosphatase activities in mitosis. Frontiers in cell and developmental biology 2018, 6, 30.

(53) Smoly, I.; Shemesh, N.; Ziv-Ukelson, M.; Ben-Zvi, A.; Yeger-Lotem, E. An asymmetrically balanced organization of kinases versus phosphatases across eukaryotes determines their distinct impacts. PLoS computational biology 2017, 13, e1005221.

(54) Grimaldi, M.; Boulahtouf, A.; Toporova, L.; Balaguer, P. Functional profiling of bisphenols for nuclear receptors. Toxicology 2019, 420, 39–45.

(55) Molangiri, A.; Varma, S.; Boga, N. S.; Das, P.; Duttaroy, A. K.; Basak, S. Gestational exposure to BPA alters the expression of glucose and lipid metabolic mediators in the placenta: Role in programming offspring for obesity. Toxicology 2024, 509, 153957.

(56) Ozden Akkaya, O.; Yağci, A.; Zik, B.; Kibria, A. G.; Güler, S.; Celik, S.; Altunbaş, K. The effect of bisphenol A on the Notch (Notch2 and Jagged2) signaling pathway in the follicular development of the neonatal rat ovary. Biotechnic & Histochemistry 2024, 99, 238–259.

(57) Chen, Z.; Zuo, X.; He, D.; Ding, S.; Xu, F.; Yang, H.; Jin, X.; Fan, Y.; Ying, L.; Tian, C.; others Long-term exposure to a ‘safe’dose of bisphenol A reduced protein acetylation in adult rat testes. Scientific reports 2017, 7, 40337.

(58) Lan, X.; Fu, L.-J.; Zhang, J.; Liu, X.-Q.; Zhang, H.-J.; Zhang, X.; Ma, M.-F.; Chen, X.M.; He, J.-L.; Li, L.-B.; others Bisphenol A exposure promotes HTR-8/SVneo cell migration and impairs mouse placentation involving upregulation of integrin-β1 and MMP-9 and stimulation of MAPK and PI3K signaling pathways. Oncotarget 2017, 8, 51507.

(59) Li, Y.; Yang, Z.; Li, W.; Xu, S.; Wang, T.; Wang, T.; Niu, M.; Zhang, S.; Jia, L.; Li, S. TOPK promotes lung cancer resistance to EGFR tyrosine kinase inhibitors by phosphorylating and activating c-Jun. Oncotarget 2016, 7, 6748.

(60) Cole, A.; Frame, S.; Cohen, P. Further evidence that the tyrosine phosphorylation of glycogen synthase kinase-3 (GSK3) in mammalian cells is an autophosphorylation event. Biochemical Journal 2004, 377, 249–255.

(61) Doble, B. W.; Woodgett, J. R. GSK-3: tricks of the trade for a multi-tasking kinase. Journal of cell science 2003, 116, 1175–1186.

(62) Smeal, T.; Binetruy, B.; Mercola, D. A.; Birrer, M.; Karin, M. Oncogenic and transcriptional cooperation with Ha-Ras requires phosphorylation of c-Jun on serines 63 and 73. Nature 1991, 354, 494–496.

(63) Patel, P.; Woodgett, J. R. Glycogen synthase kinase 3: a kinase for all pathways? Current topics in developmental biology 2017, 123, 277–302.

(64) Wang, T.; Xie, C.; Yu, P.; Fang, F.; Zhu, J.; Cheng, J.; Gu, A.; Wang, J.; Xiao, H. Involvement of insulin signaling disturbances in bisphenol A-induced Alzheimer’s diseaselike neurotoxicity. Scientific reports 2017, 7, 7497.

