## Supplementary document for "Integrated proteomic and phosphoproteomic profiling reveals mechanisms of Bisphenol-A induced placental toxicity"

**Supplementary Information**

**Supplementary File S1:** The differentially expressed proteins and their enriched Reactome pathways post BPA exposure in HTR8/SVneo cells.

**Supplementary File S2:** The mock and BPA unique phospho-proteins and their respective Reactome enriched pathways.

**Supplementary File S3:** The 75 SWATH identified kinases with their KEGG pathway enrichment. The regulated kinases were highlighted in yellow with their known targets of phosphorylation.

**Supplementary File S4:** The Reactome pathway enrichment of human and mice c-JUN target proteins.

**Supplementary Figure S1:** Distribution of differential parameters and mock phospho-protein enriched pathways.

**Supplementary Figure S2:** Exploring the identified kinases.

**Supplementary Figure S3:** c-JUN target associated pathways.

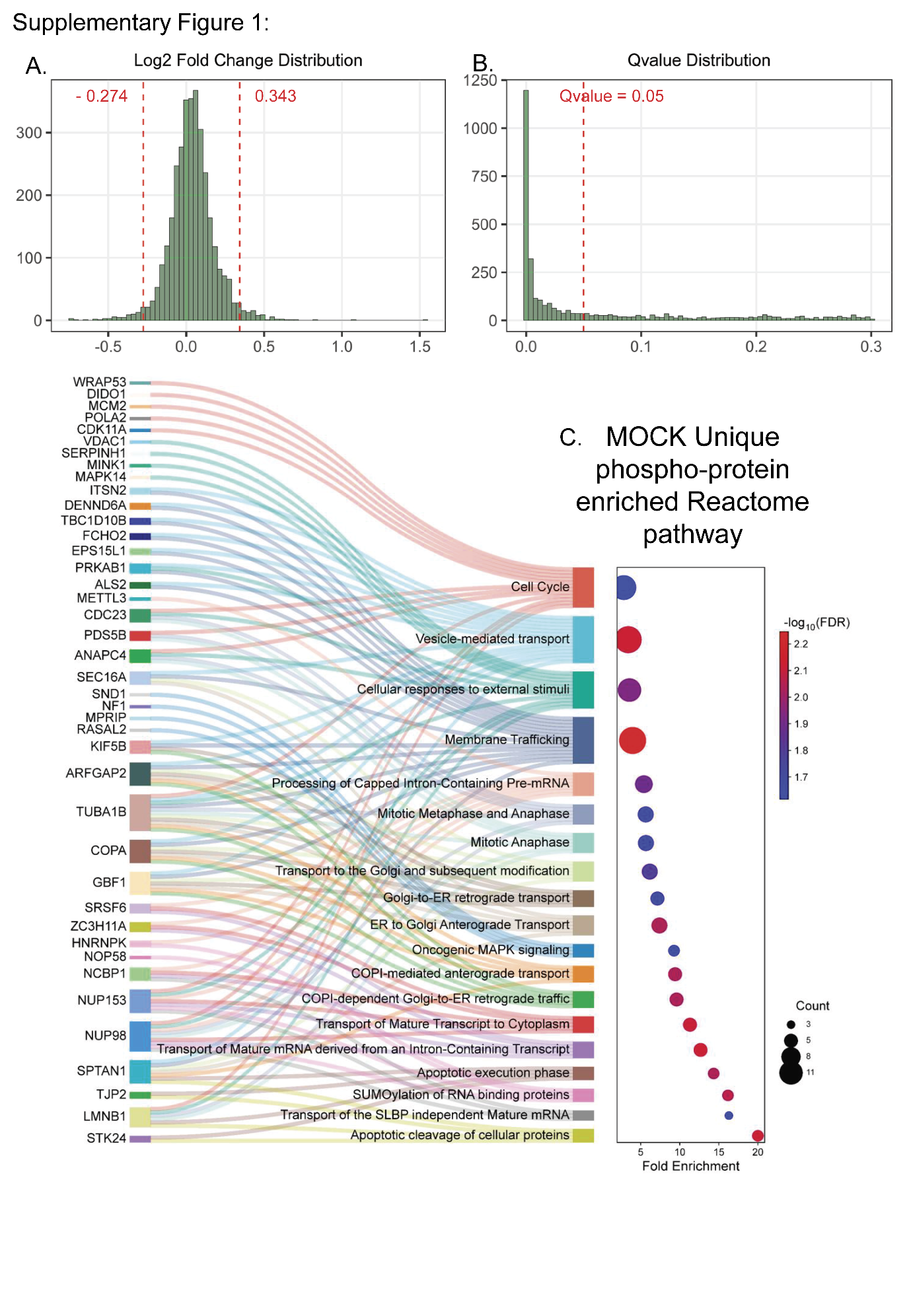

**Supplementary Figure S1: Distribution of differential parameters and mock phospho-protein enriched pathways.** The histograms display log2 fold change **(A)** and q-value **(B)** distribution of the differential proteome identified protein features. The calculated cutoffs are highlighted with red dotted lines. The mock unique phospho-protein enriched top 20 Reactome pathways are visualized by Shankey diagram **(C)**. The fold enrichment is plotted in x-axis; dot size represents protein counts and colours shows adjusted p-values.

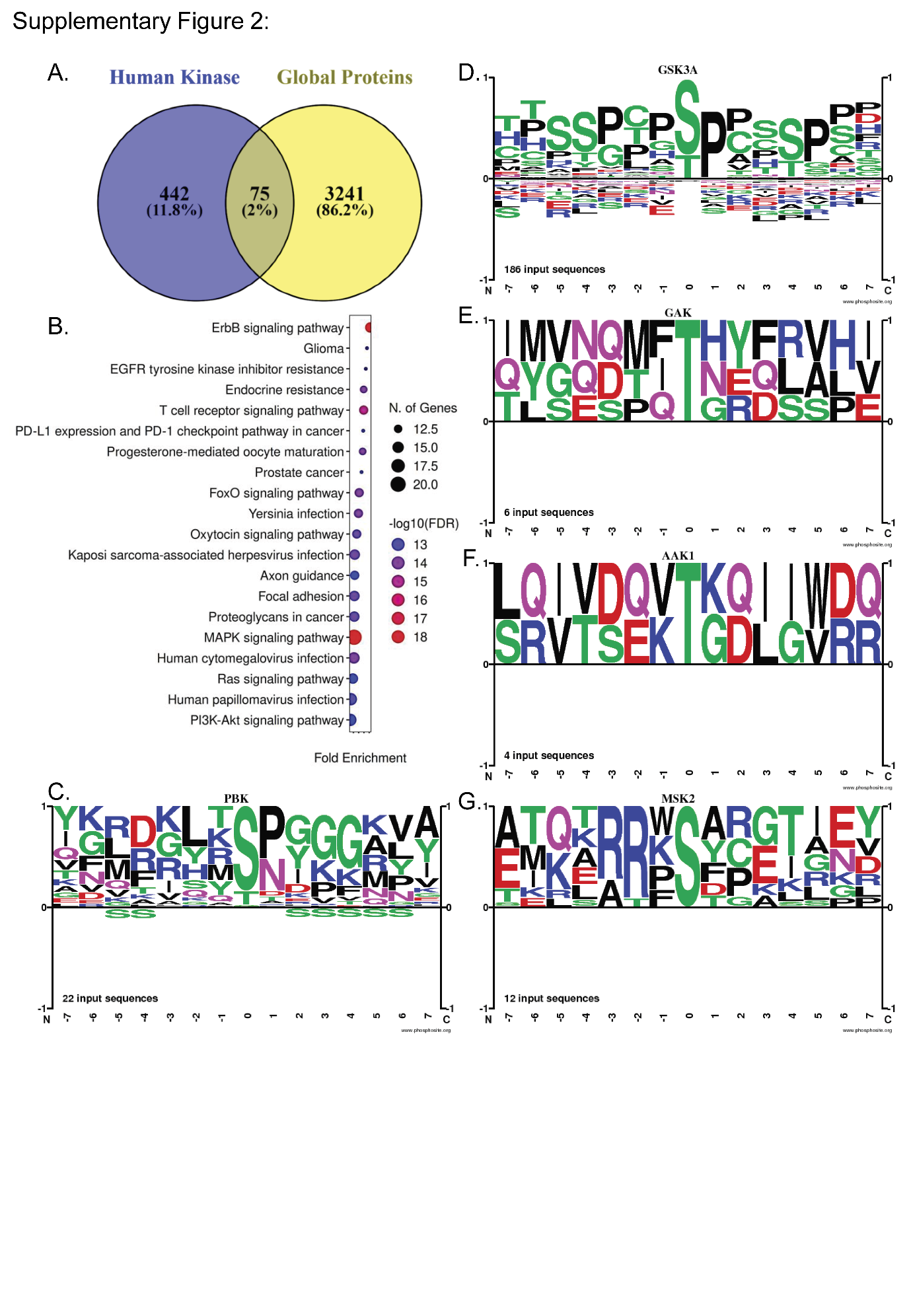

**Supplementary Figure S2: Exploring the identified kinases (A)** The Venn diagram shows overlap of know human kinases with the proteins identified in global HTR8/SVneo proteome. **(B)** The bubble plot displayed the top 20 KEGG pathways enriched by this 75 identified kinases. The dot size and colour represent protein count and adjusted p-values respectively. A sequence logo representation of conserved target motifs for PBK **(C)**, GSK3α **(D)**, GAK **(E)**, AAK1 **(F)**, and MSK2/RPS6KA4 **(G)** were visualized. The enrichment of 7 upstream and downstream amino acid residues surrounding the phosphorylation sites were plotted with positive values showing overrepresentation and negative values showing under-representation. The number of known sequences for each kinases is mentioned in the lower left corner of the plot.

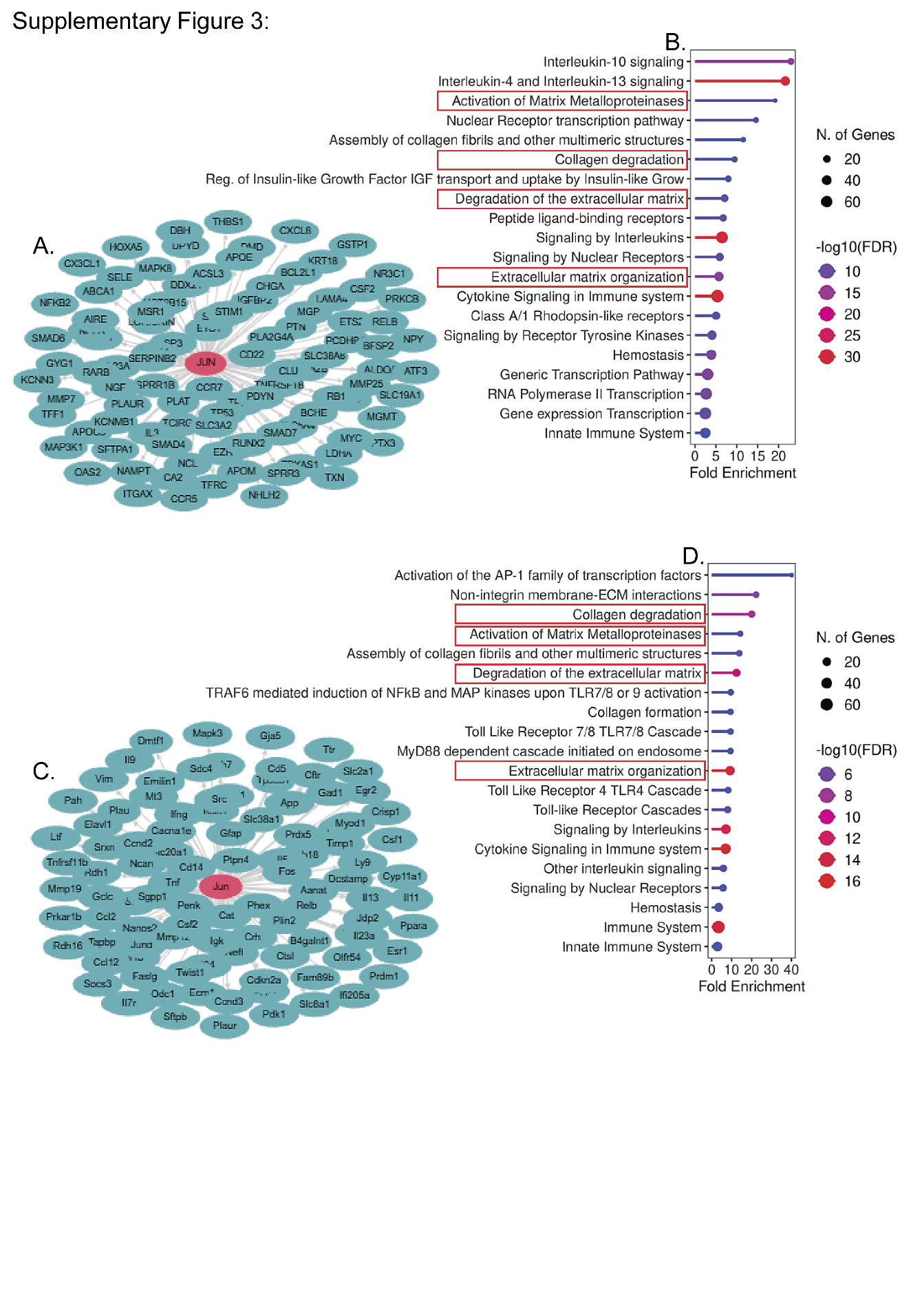

**Supplementary Figure S3: c-JUN target associated pathways.** The human **(A)** and mice **(C)** target proteins for c-JUN are visualized with a network plot where edges from each target protein is connected with red coloured c-JUN node. The lollipop plot displayed the top 20 Reactome pathways enriched by either human **(B)**, or mice **(D)** target genes. The colour and size of lollipop is represented by adjusted p-value and protein count respectively. The relevant common pathways are highlighted in red.
